# Beneficial ‘inefficiencies’ of western ranching: Flood-irrigated hay production sustains wetland systems by mimicking historic hydrologic processes

**DOI:** 10.1101/2023.12.10.571036

**Authors:** J. Patrick Donnelly, Kelsey Jensco, John S. Kimball, David Ketchum, Daniel P. Collins, David E. Naugle

**Author notes:** Corresponding author. E-mail address (J.P. Donnelly).

## Abstract

Ranching in the American West has long relied on riparian ecosystems to grow grass-hay to feed livestock in winter and during drought. Producers seasonally flood grasslands for hay production using stream diversions and low-tech flood-irrigation on riparian floodplains. Inundation mimics natural processes that sustain riparian vegetation and recharge groundwater. The recent doubling in use of more efficient irrigation approaches, such as center-pivot sprinklers, threatens to accelerate climate change impacts by unintentionally decoupling more inefficient, traditional practices that sustain riparian systems. To address this information gap, we developed an exhaustive spatial inventory of grass-hay production and combined it with monthly surface water distributions modeled from satellite data. Surface water data were classified by wetland hydroperiod and used to estimate the proportion of wetlands supported by grass-hay production in the Intermountain West, USA. Elevation and proportion of grass-hay relative to other irrigated lands were enumerated to examine differences in their positions and abundance within landscapes. Lastly, we overlaid the delineated grass-hay wetlands with LANDFIRE pre-Euro-American Settings layer to quantify the efficacy of flood irrigation in mimicking the conservation of historical riparian processes. Findings suggest that inefficient grass-hay irrigation mirrored the timing of natural hydrology, concentrating ∼93% of flooded grasslands in historical riparian ecosystems, affirming that at large scales, this ranching practice, in part, mimics floodplain processes sustaining wetlands and groundwater recharge. Despite representing only 2.5% of irrigated lands, grass-hay operations supported a majority (58%)of temporary wetlands, a rare and declining habitat for wildlife in the Intermountain West. Tolerance for colder temperatures confined grass-hay production to upper watershed reaches where higher value crops are constrained by growing degree days. This novel understanding of grass-hay agroecology highlights the vital role of working ranches in the resilience and stewardship of riparian systems.

## 1. Introduction

Agricultural ecosystems are conspicuous consumers and producers of ecosystem services (ES), harnessing environmental inputs (e.g., water and soil) to provision food and fiber, benefitting human wellbeing. Growing recognition that agriculture is capable of broader ES supporting environmental regulation (e.g., water quality) and function (e.g., soil formation) is increasing its value as a mechanism for ecological sustainability (MEA 2005; Power, 2010; Swinton et al., 2007). Predicted impacts of climate change are expected to influence agricultural ES and food security, particularly irrigated agriculture (i.e., human application of water to cultivate crops), which comprises one-fifth of the cropped area globally but produces 45% of the world’s food (Döll and Siebert, 2002). Increased frequency and severity of extreme drought events are anticipated to accelerate water scarcity, depleting yields through curtailment of agricultural irrigation and growing competition from expanding urban and industrial demands (Gornall et al., 2010). Addressing emerging gaps in water resources will require an increased understanding of agricultural ES and their ability to bolster climate resilience through ecologically sustainable irrigation practices.

Crop production in the Intermountain West, USA (Fig. 1) is dependent on irrigation, with agriculture consuming over 80% of extracted freshwater (Dieter, 2018). Government investment in large-scale reclamation projects has enabled the capture and conveyance of surface water, supporting agriculture, industry, and urbanization in otherwise water-limited landscapes (Hansen et al., 2009). Stream runoff from mountain snowpack, stored behind large reservoirs, has historically provided reliable water sources to downstream users, decoupling crop yields from historic climatic constraints. Emerging climate change effects and increasing water demands are now outpacing the functional capacity of reclamation infrastructure to offset drought (Siirila-Woodburn et al. 2021). For example, the overallocation of water resources under a continuing 22-year drying trend in the Colorado River basin has reduced Lake Mead, the largest reservoir in the U.S., to 27% of capacity in 2022, increasing demands for conservation measures that offset diminished water supply through more efficient agricultural use (Richter et al., 2017).

**Fig. 1.**
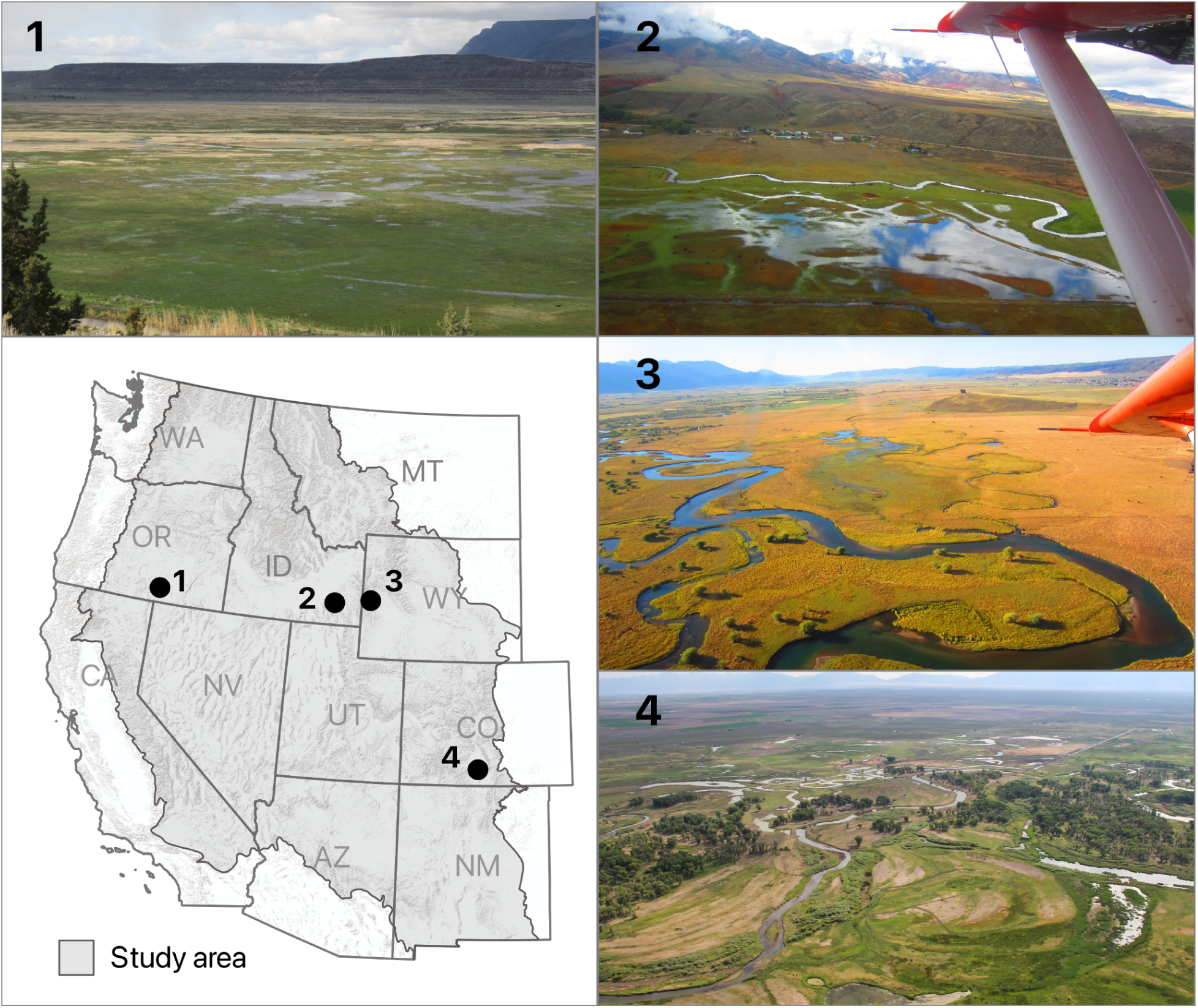
Study area map defined by the Intermountain West, USA. Images characterize agroecological settings associated with regional flood-irrigated grass-hay production: 1) Warner Valley, Oregon; 2) Marsh Valley, Idaho; 3) Star Valley, Wyoming; 4) San Luis Valley, Colorado.

Riparian floodplains and wetlands account for only 1-3% of the Intermountain West’s geography (Tiner, 2003). While most have been significantly altered through water extraction for agricultural and urban demands, riparian ecosystems remain fundamental drivers of hydrologic cycles crucial to structuring biodiversity and landscape ecosystem function (Patten, 1998). Since Euro-American settlement, ranching has relied on riparian ecosystems for fertile floodplain soils and irrigation supporting grass-hay production needed for livestock (Talbert et al., 2007). Flood-irrigation sustaining this practice results in seasonally flooded grasslands linked to contemporary ES through local studies that identify associated groundwater recharge as a key component of riparian ecosystem resilience that maintains wetland function, cools water temperatures, and bolsters late summer flows through in-stream groundwater discharge (Alger et al., 2021; Essaid and Caldwell, 2017; Ferencz and Tidwell, 2022; Gordon et al., 2020; Welsh et al., 2013). Integrated over large basins, return flows from flood-irrigation (generated through groundwater recharge and subsequent in-stream discharge) can provide substantial quantities of additional in-stream baseflow. For example, a regional study conducted by Lonsdale et al. (2020) found that of 12.6 km3 (10.5 million-acre feet) that are diverted annually for irrigation in areas of Montana (where flood-irrigation is the primary irrigation practice), 9.86 km3 (8.0 million-acre feet) go un-consumed by evapotranspiration and likely contribute to groundwater recharge.

Climate change strategies meant to offset increasing water scarcity through the adoption of more efficient irrigation infrastructure (e.g., pressurized pipes and center pivot sprinklers) frequently view grass-hay flood-irrigation as wasteful without additional consideration of lost ES that may occur as a result of conversion (Richter et al., 2017; Scott et al., 2014). Our limited understanding of ES provided from flood-irrigated grass-hay production (hereafter “grass-hay production”) could speed up environmental degradation by unintentionally decoupling agricultural irrigation practices that contribute to the sustainability of riparian ecosystems. To fill this information gap, we developed an exhaustive spatial inventory of grass-hay production in the Intermountain West. Satellite data was used to model monthly (Jan-Dec) surface water extent averaged from 2013 to 2022. Results were classified by wetland hydroperiod and used as indicators for ES to determine the proportion of wetland resources (i.e., flooded grasslands) supported by grass-hay production in the Intermountain West. We calculated the elevational distributions and proportion of grass-hay production relative to other irrigated lands to explore differences in their specific positions and abundance within landscapes. Lastly, grass-hay wetlands were compared with pre-Euro-America “ecological systems maps” to assess their overlap with historical riparian ecosystems and potential for supporting groundwater recharge. Study outcomes provide the first-ever spatially explicit framework linking ranching and grass-hay production to ES supporting riparian ecosystem function in the Intermountain West.

## 2. Materials and Methods

### 2.1 Study area

The study area encompassed the Intermountain West region of the United States (Fig. 1), that included areas between the front ranges of the Rocky Mountains on the east and the Cascade Mountains and Sierra Nevada (Mountains) on the west. The region consists of extensive mountains and plateaus separated by broad valleys and lowlands. The climate is arid to semi-arid, with marked seasonal temperature extremes consisting of cold, wet winters and warm to hot, dry summers. Aridity is the result of rain shadows caused by mountain ranges intercepting wet air masses that concentrate winter precipitation as snowpack in upper elevations. Annual precipitation is highly variable, controlled by changing ocean surface temperatures that shape global atmospheric circulation patterns impacting the region (Rajagopalan and Lall, 1998). Merged Level-3 North American ecoregions defined the study area boundary (Wiken et al., 2011).

Hydrology, driven primarily by snowmelt and runoff, concentrates surface water and wetlands in riparian floodplains, where seasonal flooding during spring and early summer supports groundwater recharge. Most wetlands occur naturally but may also include publicly managed wetlands that are intentionally flooded for the benefit of wildlife (hereafter “natural wetlands”). Seasonally flooded grasslands associated with cultivated grass-hay are important agroecological wetland resources on private working ranches (sensu Downard and Endter-Wada, 2013).

Outside large urban centers, the region is primarily rural, supporting agriculture, natural resource extraction (e.g., energy, timber, and mining), and outdoor recreation industries. Forested mountains, rangelands, and cultivated agriculture are the dominant land-cover types. Aridity requires nearly all crops to be irrigated. Public lands owned primarily by the U.S. Forest Service and the U.S. Bureau of Land Management make up 63% of the study area (USGS 2018), supporting extensive private livestock grazing operations that seasonally rotate cattle and sheep between government and privately owned rangelands to meet forage needs (Fleischner, 1994).

### 2.2 Grass-hay delineations

Areas of grass-hay production were delineated as digitized polygons in a Geographical Information System (GIS) using on-screen image interpretation. Spatial analysts, familiar with regional agroecology, employed direct and indirect interpretation techniques by overlaying ancillary land-use, land-ownership, and land-cover datasets (Table 1) onto recently acquired (2013-22) true-color, high-resolution (≤ 1-meter) satellite imagery (sensu Lillesand et al., 2015). For example, irrigation ditches in grass-hay fields visible in optical satellite imagery were used as an indirect determinant of flood-irrigation practices. Detections were aided by color-infrared and normalized difference vegetation indices (NDVI) derived from multispectral satellite imagery, making it possible to visualize variation in herbaceous productivity (Pettorelli et al., 2005). Aridity in the region promoted high contrast between irrigated and non-irrigated vegetation, aiding the detection of grass-hay production. Additional inference to indirect detection measures was provided by overlaying herbaceous productivity (i.e., NDVI greenness) information with land-cover classification maps depicting perennial grass cover (Allred et al., 2021). Accuracy assessments from ancillary land-cover datasets were reviewed by analysts to aid interpretation and reduce the effects of compounding error from low-confidence data.

**Table 1.** Ancillary datasets used in support of flood-irrigated grass-hay delineations.

| Variable | Source |
| --- | --- |
| Land-cover |  |
| Croplands Data Layer | (NASS 2019) |
| Rangeland Analysis Platform - Perineal Grass cover | (Allred et al., 2021) |
| Land-Use Land-ownership |  |
| Common land Unit Dataset | (NRCS 2010) |
| IrrMapper - irrigated lands | (Ketchum et al., 2020) |
| Protected Areas Database | (USGS 2018) |
| Satellite Imagery |  |
| Google Imagery, Natural color ≤ 1-meter resolution | (Google, 2022) |
| Sentinel 2, Color infrared 10-meter resolution | (European Space Agency, 2015) |
| Sentinel 2, NDVI 10-meter resolution | (European Space Agency, 2015) |
| Geomorphic |  |
| Slope 10-meter resolution | (USGS 2012) |

The accuracy of grass-hay delineations was assessed using a stratified binary sampling approach by randomly generating 5,000 sample points distributed evenly inside and outside of grass-hay polygons. Random points generated outside grass-hay polygons were restricted to other irrigated lands as calculated from mean annual conditions for the study period (2013-22) using the IrrMapper Irrigated Lands dataset (Ketchum et al., 2020). Irrigated derived lands from IrrMapper included all agriculture in addition to urban uses limited to grass turf for golf courses and large athletic fields that made up a minor component of overall estimates. A binary classification was applied to each point, identifying the presence or absence of grass-hay production using an independent analyst and on-screen image interpretation methods previously described. Results were assessed using Cohen’s Kappa statistic (Cohen, 1960) to estimate overall data accuracy.

### 2.3 Wetland surface water hydrology

Monthly surface water hydrology was modeled to determine the proportion and types of wetlands supported by grass-hay production. Following methods outlined by Donnelly et al. (2021), surface water measurements were derived from Landsat 8 and 9 Operational Land Imager satellite imagery using a continuous 30x30 meter pixel grid. Monthly (Jan-Dec) estimates represented mean surface water conditions from 2013 to 2022. This approach made it possible to represent recent ecological conditions while reducing the influence of climate-induced outliers occurring during the study period. Using standards similar to Cowardin et al. (1979), surface water conditions were classified into hydroperiods by totaling the months individual pixels were inundated from January to December. Hydroperiod is a key delimiter of wetland function important to determining ecological values (sensu Minckley et al., 2013). Classes included “temporary” (flooded < 2 months), “seasonal” (flooded > 2 and < 9 months), and “semi-permanent” (flooded > 9 months). Large deep water bodies (e.g., reservoirs) were omitted from our analysis to remove bias from surface water features unassociated with most riparian ecosystems in the Intermountain West. Detailed descriptions of remote sensing methods used to estimate surface water conditions are provided as supplemental materials (*see* Supplementary material, Remote Sensing Methods, Surface water modeling).

### 2.4 Ecosystem service indicators

Wetland hydroperiod classes (i.e., temporary, seasonal, and semi-permanent) were summarized within grass-hay polygons as indicators of ES supporting wetlands and riparian ecosystem function (e.g., groundwater recharge). Area measures were summarized by month (Jan-Dec) as a time series hydrograph. Procedures were repeated for natural (i.e., non-grass hay) wetlands. Overall grass-hay and natural wetland abundance was estimated at an annual scale by summing monthly area measures by hydroperiod classes.

We estimated the proportion of irrigated agriculture supporting grass-hay wetlands in the Intermountain West using IrrMApper Irrigated Lands dataset. Calculations were made by dividing the area of grass-hay by the total area of irrigated land. The area measurement used for grass-hay wetlands was calculated as a maximum surface water footprint derived from the spatial aggregation of monthly estimates (Jan-Dec). This approach excluded portions of inventoried grass-hay production (i.e., polygons) that our analysis determined were unsupportive of measurable grassland flooding associated with wetland ES. Results were summarized by state to facilitate the integration of our results into regional planning and conservation strategies. All area measures were representative of mean conditions from 2013 to 2022.

### 2.4 Historic ecosystems

Historical ecological settings underlying grass-hay production were identified by intersecting grass-hay wetlands with LANDFIRE’s Biophysical Settings layer ー an ecological systems classification representing landscape conditions from pre-Euro-American settlement (USDA 2018). This process made it possible to quantify overlap with historical riparian ecosystems (defined using NatureServe’s terrestrial ecological systems of the United States - natureserve.org) and grass-hay wetlands supporting contemporary riparian ES. Overlap was summarized by aggregating “riparian ecological system classes” into a single riparian class (Table 1). The aggregated riparian layer was processed using a majority single-pass filter (90-meter radius) to increase the spatial heterogeneity of the data (Cardille et al., 2023). This approach enhanced the prevalence of dominant classes, providing a more uniform representation of riparian ecosystems. (*see example* Fig. S1). The maximum footprint of grass-hay wetlands was used to determine overlap with historical riparian areas. Proximity of grass-hay wetlands occurring outside historic riparian ecosystems was estimated using a per-pixel Euclidean distance function. Results were reported as a global distance mean. To estimate the overlap between historical riparian areas and other (i.e., non-grass-hay) irrigated lands, the aggregated riparian layer was intersected with IrrMapper data using mean conditions from 2013 to 2022. Areas of IrrMapper data overlapping grass-hay wetlands were removed before the analysis to prevent double counting.

**Table 1a.**
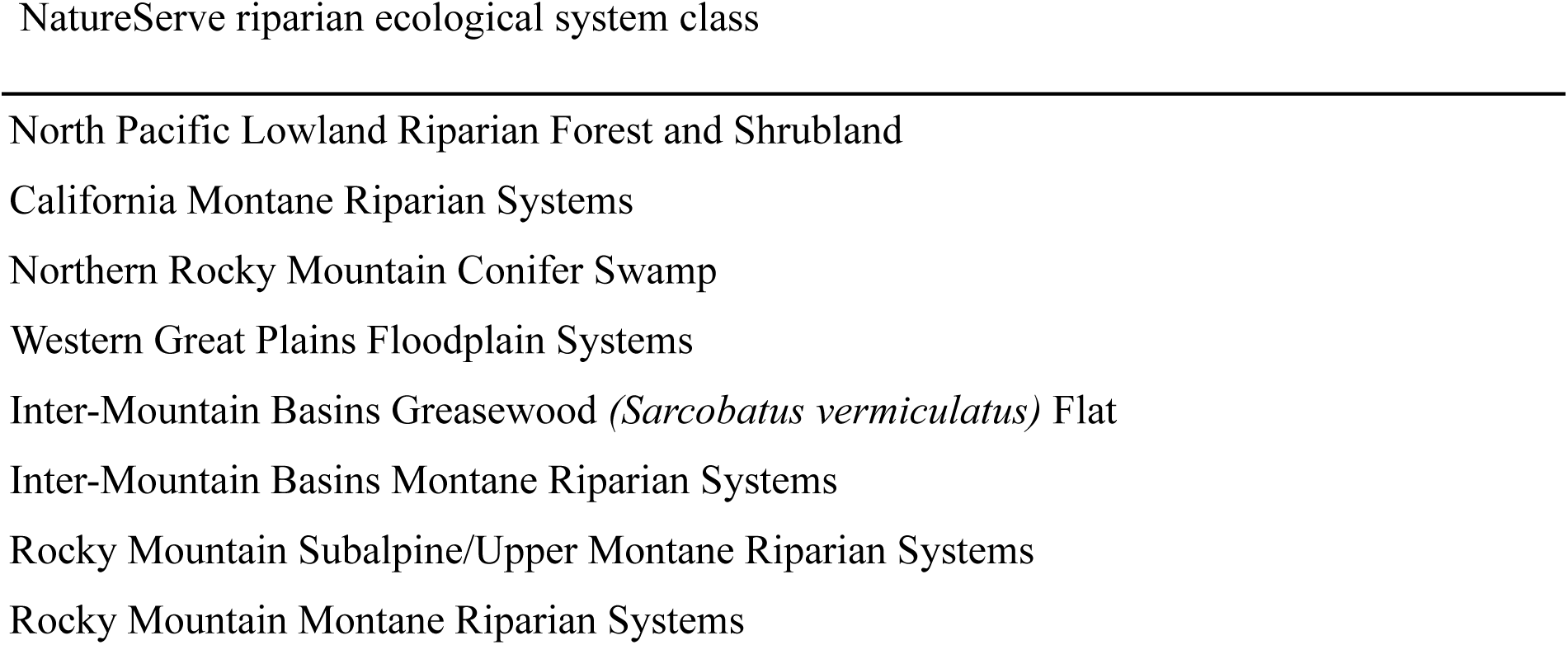
NatureServe’s terrestrial ecological system classes used for estimation of pre-Euro-American riparian ecosystem extent.

### 2.5 Elevation

Lastly, we calculated the elevations of grass-hay production and other irrigated lands to compare differences in landscape position. Elevation measures were derived by spatially intersecting the maximum footprint of grass-hay wetlands with the U.S. Geological Survey 10-m National Elevation Dataset (USGS 2012). This process was repeated for other irrigated lands using the previously described IrrMapper dataset (mean conditions from 2013-2022). Elevational distributions were summarized as boxplots.

### 2.6 Data processing

All image processing and raster-based analyses were conducted using Google Earth Engine, a cloud-based geospatial processing platform (Gorelick et al., 2017). All GIS analyses were performed using QGIS (QGIS Development Team, 2020). Plotting and statistical analyses were generated using the R environment (R Core Team, 2019; RStudio Team, 2019), including R-package Tidyverse (Wickham et al., 2019).

## 3. Results

Annually, grass-hay production accounted for 57.8%, 19.9%, and 6.1% of temporary, seasonal, and semi-permanent wetlands in the Intermountain West from 2013 to 2022. At broad spatial and temporal scales, grass-hay wetlands exhibited monthly trends similar to natural wetlands but differed in relative abundance (Fig. 2). A majority of temporary wetlands were attributed to grass-hay production from April through July, with the June peak accounting for over two-thirds (68.1%) of availability (Table 2). Seasonal wetlands followed similar trends, with annual peaks occurring from May to June. Surface water trends for semi-permanent wetlands remained relatively constant for grass-hay and natural wetlands, with declines in January and December likely due to freezing and snow cover.

**Fig. 2.**
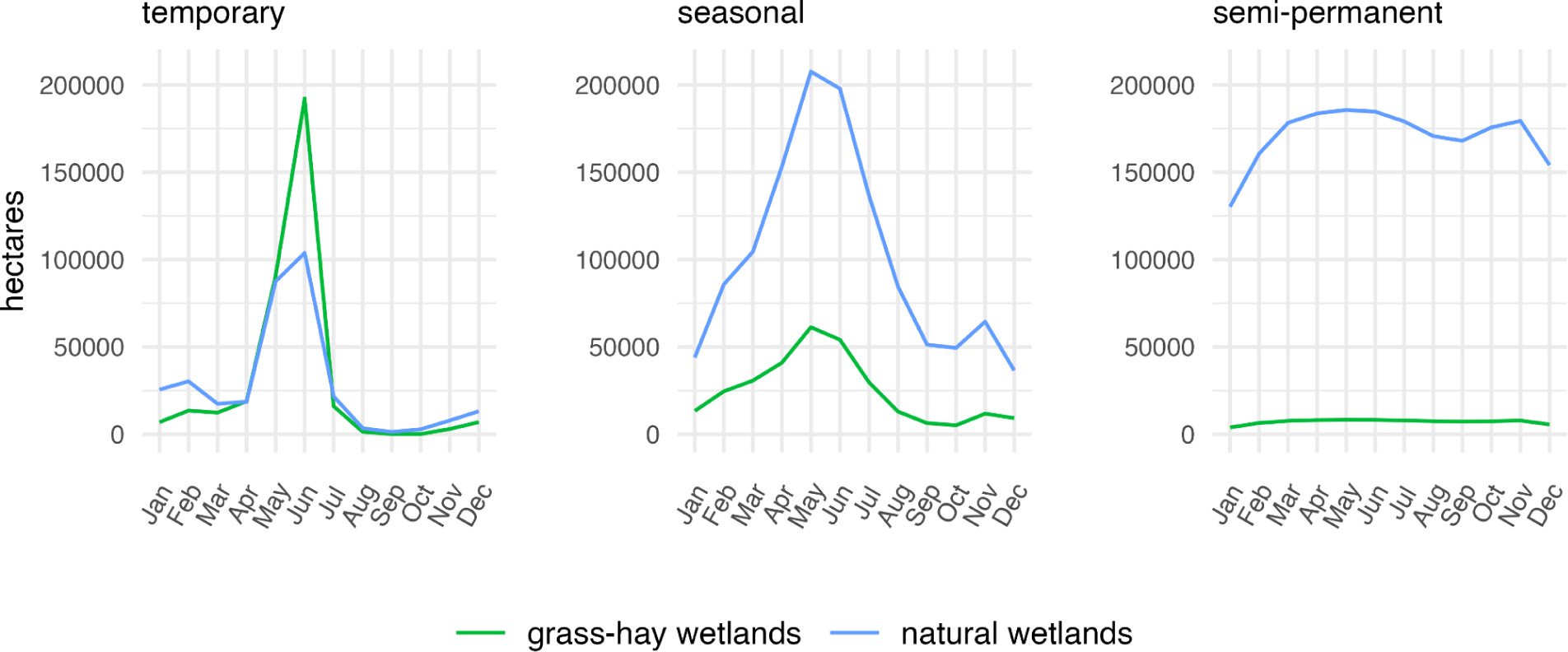
Hydrographs depicting mean monthly surface water abundance by wetland hydroperiod class (temporary, seasonal, and semi-permanent). Measures shown for flood-irrigated grass hay wetlands and natural wetlands in the Intermountain West as mean conditions from 2013 to 2022. Hydrographs for individual states are available as Supplementary Material (*see* Figures S2-S10).

**Table 2.** Monthly proportion of wetland hydroperiod classes supported by flood-irrigated grass-hay production in the Intermountain West (2013-22).

| Hydroperiod | Jan | Feb | Mar | Apr | May | Jun | Jul | Aug | Sep | Oct | Nov | Dec |
| --- | --- | --- | --- | --- | --- | --- | --- | --- | --- | --- | --- | --- |
| temporary | 34.9% | 38.9% | 44.8% | 54.3% | 55.8% | 68.0% | 54.5% | 36.0% | 16.4% | 5.8% | 29.7% | 41.1% |
| seasonal | 23.3% | 22.4% | 22.8% | 21.2% | 22.9% | 21.6% | 18.0% | 13.5% | 11.2% | 9.5% | 15.5% | 20.3% |
| semi-perm | 4.2% | 5.7% | 6.1% | 6.3% | 6.4% | 6.4% | 6.4% | 6.5% | 6.4% | 6.2% | 6.3% | 5.1% |

**Table 3.** Intermountain West occurrence of flood-irrigated grass-hay wetlands and total irrigated land by state. All measures are shown in hectares.

| State | Grass-hay wetland area by state | Proportion of total grass-hay wetland area by state | Total land irrigated by state | Grass-hay wetland as proportion of lands irrigated by state |
| --- | --- | --- | --- | --- |
| Arizona | 0 | 0.0% | 31,295 | 0.0% |
| New Mexico | 0 | 0.0% | 825,967 | 0.0% |
| Washington | 8,267 | 2.5% | 2,138,274 | 0.4% |
| California | 11,744 | 3.5% | 562,953 | 2.1% |
| Utah | 18,922 | 5.7% | 934,520 | 2.0% |
| Nevada | 20,886 | 6.3% | 607,572 | 3.4% |
| Colorado | 31,563 | 9.5% | 1,122,732 | 2.8% |
| Oregon | 46,626 | 14.1% | 1,488,584 | 3.1% |
| Montana | 56,843 | 17.2% | 1,061,181 | 5.4% |
| Idaho | 62,518 | 18.9% | 3,606,640 | 1.7% |
| Wyoming | 73,734 | 22.3% | 1,034,691 | 7.1% |

Grass-hay wetlands accounted for 2.5% (∼330,000 ha) of irrigated lands (13.4 million ha) in the Intermountain West (Fig. 3). The largest concentrations occurred in Wyoming, Idaho, Montana, Oregon, and Colorado, which combined accounted for over 80% of abundance (Table 2). Non-grass-hay irrigated agriculture was most abundant in Washington and Idaho, resulting in relatively low irrigated land-to-grass-hay wetland proportions. Irrigated proportions were highest in Wyoming and Montana, where grass-hay wetlands accounted for 7.1% and 5.4% of irrigated lands. Limited grass-hay production in Arizona and New Mexico lacked the flood-irrigation practices needed to support wetland conditions.

**Fig. 3.**
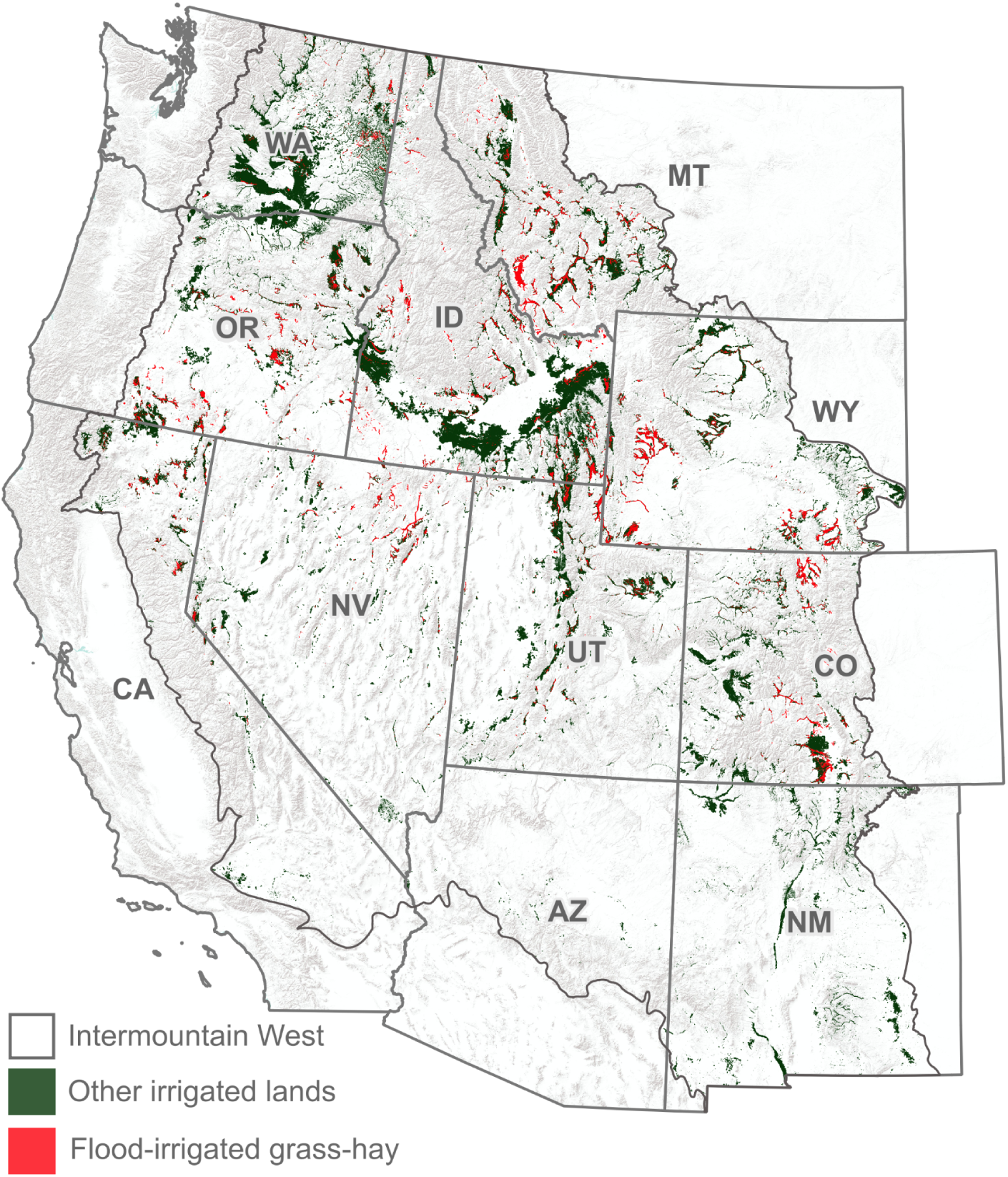
Intermountain West irrigated agriculture shown as flood-irrigated grass-hay and other irrigated lands. Delineations represent mean conditions from 2013 to 2022.

Pre-Euro-American riparian ecosystems encompassed 92.8% of grass-hay wetlands in the Intermountain West (Fig.4). Omitted areas (∼24,000 ha) were proximal with per-pixel distances averaging 188 meters from historical riparian sites. Only 11.8% of non-grass-hay irrigated lands overlapped historic riparian ecosystems. Grass-hay production occurred at median elevations approximately 500 meters higher than other irrigated lands (Fig. 5). While the lower quartile range of grass-hay production (1368 m) overlapped the upper range of other irrigated crops (1583 m), distributions characterized broad-scale patterns of landscape position that place grass-hay at notably higher elevations. The mapping accuracy of cultivated grass-hay practices was high, with an estimated Cohen’s Kappa statistic of 0.99. During evaluations, minor rates of omission errors were detected in Oregon and Colorado. Corrections were appended to the dataset and incorporated in the final analysis.

**Fig. 4.**
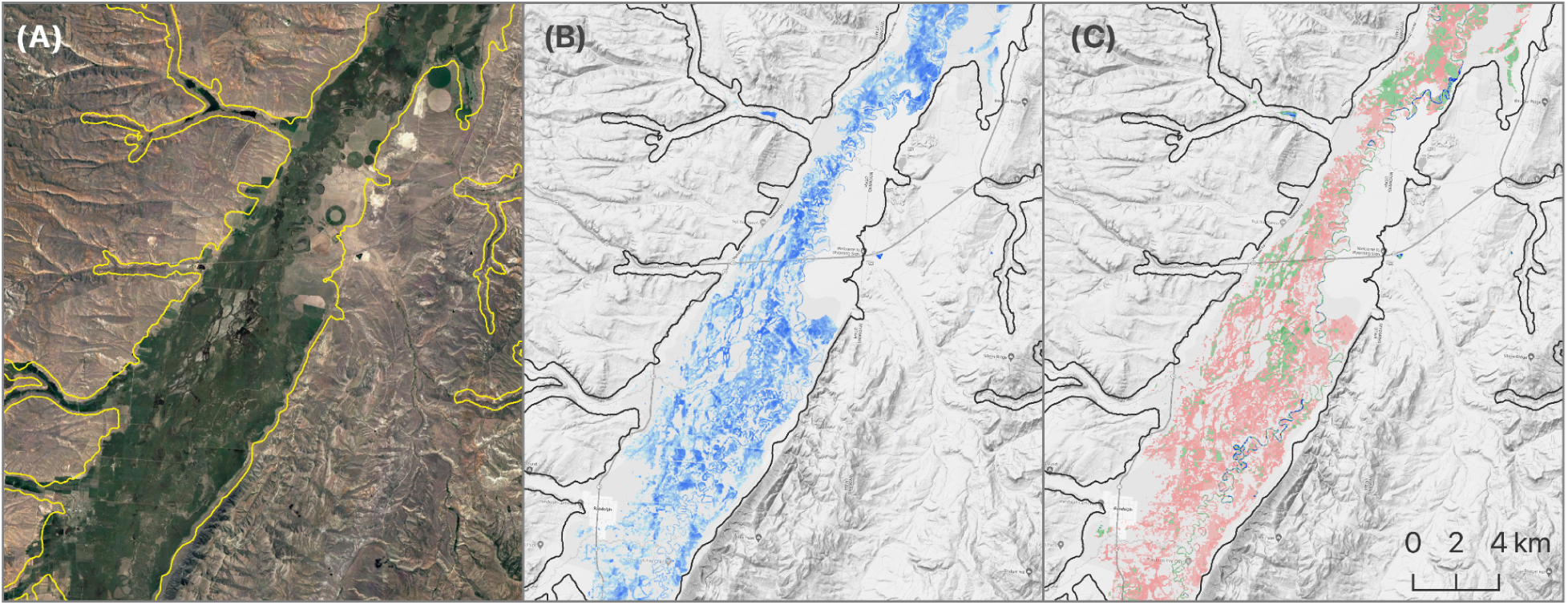
Pre-Euro-American riparian ecosystem extent (yellow/black polygon; A, B, C) estimated from LANDFIRE’s Biophysical Settings layer for area of the upper Bear River, Utah. Satellite image (A) shows dark green extent of irrigated agricultural, primarily flood-irrigated grass-hay. Satellite-based surface water delineation shown in blue (B) depicts the extent of grass hay flood-irrigation (mean conditions for June 2013-22). Lighter blue shades represent lower per-pixel proportions of surface water measured (see Supplementary Materials, Remote sensing methods, Surface water modeling). Surface water is shown by wetland hydroperiod class (C) as temporary (pink), seasonal (green), and semi-permanent (blue).

**Fig. 5.**
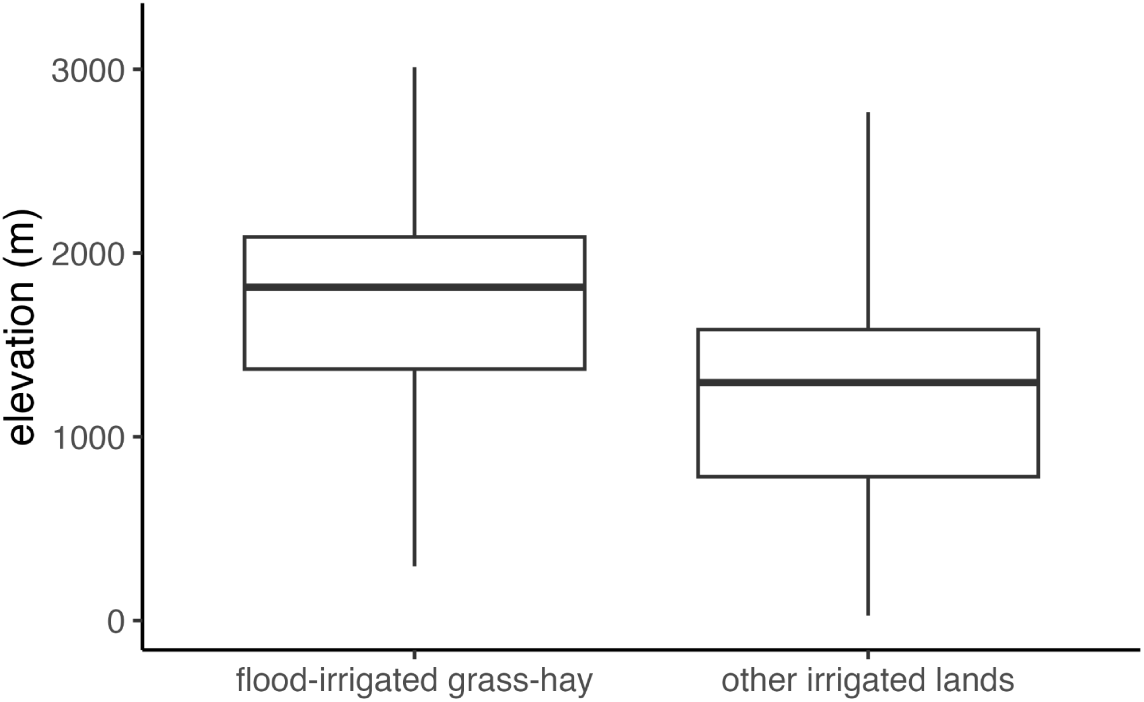
Elevational distribution of flood-irrigated grass-hay wetlands and other irrigated lands in the Intermountain West. Boxes represent the interquartile range (IQR); the median value is the line dividing the box horizontally; and whiskers, 1.5 times the IQR.

## 4. Discussion

Ours is the first analysis to quantify wetland and riparian ES linked to grass-hay production in the Intermountain West. Findings fill a crucial knowledge gap in the role that ranching plays in mimicking historic, ecosystem-scale hydrologic processes that have been lost to more efficient water uses elsewhere. Although grass-hay wetlands represented only 2.5% of irrigated lands, they accounted for 68.1% of temporary wetlands and 22.9% of seasonal wetland resources during the peak May-June irrigation season. Irrigation concentrated 92.8% of flooded grasslands in historical riparian ecosystems that mirrored the timing of natural hydrology, affirming this practice, in part, mimics floodplain processes sustaining wetlands and groundwater recharge. This improved understanding of grass-hay agroecology elevates the urgency to maintain and strategically conserve these “beneficial inefficiencies” on western working lands.

Casual linkages between grass-hay ES and riparian ecosystems were supported through detailed, evidence-based studies that connected our results to local-scale outcomes. For example, analyses by Essaid and Caldwell (2017) tied flood-irrigation supporting grass-hay production in the upper Smith River watershed, Montana, to increased in-stream groundwater discharge, leading to cooler water temperatures and improved thermal conditions for wild trout populations during summer. More recent work examining grass-hay production in the Upper Green River Basin, Wyoming, attributed over two-thirds of flood-irrigation water use to ES, including groundwater recharge, late summer in stream discharge, and consumption (i.e., evapotranspiration) supporting native riparian vegetation. (Gordon et al., 2020). We acknowledge that local ecological complexities can influence relationships between irrigation practices and ES benefits through variance in the mechanistic drivers underlying hydrologic function (i.e., geology, climate, and land use). For example, Kendy and Bredehoeft (2006) identified relationships between higher stream flows and agricultural irrigation in the Gallatin River Valley, Montana; however, lag effects caused by underlying hydrologic conditions delayed in-stream groundwater discharge until winter, when ecological benefits for aquatic systems (i.e., cooler water temperatures) were limited.

Trends between grass-hay and natural wetlands were aligned through shared hydrologic dependence on riparian ecosystems. While this study did not quantify irrigation sources directly, high concentrations of grass-hay wetlands (92.8%) within historic riparian sites suggest irrigation reliance on local surface water diversions (sensu BHWC 2023; Essaid and Caldwell, 2017; Lonsdale et al., 2020). We speculate through our results that natural wetland flooding was induced by spring snowmelt and high stream flows that concurrently supported off-channel diversions, directing water into irrigation ditches and onto adjacent agricultural floodplains. Limited grass-hay contributions to “seasonal” (19.9%) and “semi-permanent” (6.1%) wetland abundance were attributed to short-duration irrigation practices and reduced water availability during summer that favored temporary wetland hydrologies. Casual observations of our data showed that in grass-hay systems, seasonal and semi-permanent wetlands were typically confined to small areas of low topographic relief often associated with abandoned stream channels or stock ponds (*see example* Fig. S11).

Model depictions attributing over half of the temporary wetlands (57.8%) in the Intermountain West to grass-hay production suggest this practice functions as part of a broader ecological network structuring wetlands and wildlife distributions in water-limited landscapes. Recent work by Donnelly et al. (in press) found that core greater sandhill crane (*Antigone canadensis tabida*) breeding areas overlaid 93% of grass-hay production in the Intermountain West, accounting for more than 60% of temporary wetland habitats supporting birds. Moreover, healthy riparian groundwater systems sustained by grass-hay production are supportive of drought-tolerant phreatophytic plant communities (Gordon et al., 2020) that bolster vegetative productivity in later summer, providing wildlife with predictable foraging opportunities during periods of seasonal drought when most upland vegetation is desiccated (Silverman et al., 2019). Studies examining greater sage-grouse *(Centrocercus urophasianus)* identify drought tolerance in grass-hay agricultural systems as a central habitat component in the bird’s life history that sustains populations during periods of prolonged drought when foraging resources are otherwise scarce (Donnelly et al., 2018). Other wildlife benefits include contributions to continental migration networks critical as wetland stopover habitats for millions of North American waterfowl moving between northern breeding and southern wintering grounds (Fleskes and Yee, 2007; Miller et al., 2005).

### 4.1 Emerging threats

Riparian ecosystems’ significant dependence on agricultural water use, highlighted here in, identifies a previously unquantified risk to these important habitats. Climate-induced trends toward increasing water scarcity in the American West have generated a need to maintain food security through improved irrigation efficiency driven by government incentive programs that encourage reduced water use (Wallander and Hand, 2011). Adopting efficient irrigation methods, such as center-pivot sprinklers, allows producers to achieve greater crop yields while reducing water consumption by delivering only the water needed to maintain crops and eliminating surplus losses from deep soil infiltration associated with flood irrigation. Between 1984 and 2018, high-efficiency irrigation practices in the Western U.S. nearly doubled, expanding from 36% to 76% of irrigated lands (Hrozencik and Aillery, 2021). While necessary to offset intensifying drought and water supply shortfalls, adaptation strategies risk unintentionally accelerating climate change impacts if consideration is not given to preserving beneficial flood-irrigation practices that are partly responsible for sustaining riparian ecosystems.

With greater tolerance for colder temperatures, grass-hay production was concentrated in upper elevations where higher-value crops are constrained by limited growing degree days. Climate projections for regions of the western U.S. suggest that by 2070, average temperatures will be 1.8°C to 3.1°C above the historical baseline of 1970-1999 (Melillo et al., 2014). Projected climate trends elevate risks to grass-hay ES through producer adoption of alternative agricultural practices and crops. With significant increases in growing degree days already documented in parts of the Intermountain West (McGuire et al., 2012), warming temperatures in the coming decades are projected to significantly expand agricultural climate zones, potentially shifting cropping patterns to higher altitudes (King et al., 2018). The emerging feasibility of higher profit cropping practices provides economic incentives for grass-hay conversion, including investments in efficient irrigation infrastructure, allowing producers to increase yields by applying water savings to the expansion of irrigated lands (Wallander and Hand, 2011).

Maintaining grass-hay ES will be predicated partly on continued water use equity among agriculture and ecological demands. Aquatic ecosystems can benefit from irrigation-induced groundwater discharge to seasonally bolster stream flows, but excessive agricultural diversions may dampen benefits if flows fall below the levels needed to sustain fish and invertebrate communities (Gordon et al., 2020). For example, Robertson and Dey (2009) have linked excessive grass-hay irrigation diversions to depleted stream flows and declining aquatic productivity in Upper Green River, Wyoming. Additionally, increasing temperatures in the Upper Colorado River Basin are predicted to decrease stream discharge by over 9% per degree Celsius of warming, reducing surface water availability for riparian and agriculture systems (Milly and Dunne, 2020). Collaborative solutions in the Upper Big Hole watershed, Montana, have demonstrated potential solutions to irrigation and aquatic sustainability that rely on volunteer and incentive-based flexibility in producer diversions that maintain viable stream flows and water temperatures (BHWC 2023).

The reallocation of agricultural water rights to meet growing urban and industrial demands may also increase as climate-induced water scarcity intensifies, threatening the sustainability of grass-hay ES. Recent water shortages in the Colorado River basin, for example, have sparked speculation from investment firms that have begun to purchase agricultural irrigation rights, betting that projected scarcity will provide a lucrative market for selling water supplies to the highest bidder (Ryder, 2021; Tracy et al., 2023). Estimates suggest that the monetary value of transferring agricultural water rights may be twice that of continued crop production (Dozier et al., 2017). Efforts in Colorado and Nevada have proposed agricultural fallowing by purchasing and repurposing rural irrigation water for municipal use (Thorvaldson and Pritchett, 2006; Welsh and Endter-Wada, 2017). Such scenarios frequently require out-of-basin water transfers, altering natural water cycles and impacting the integrity of riparian ecosystems (Zhuang, 2016). Additionally, loss of irrigation can increase subdivision risk that removes wildlife-compatible land-use practices in rural landscapes as producers sell off land for development following its reduced agricultural valuations (Dozier et al., 2017).

### 4.2 Conclusions

Grass-hay production has garnered limited attention for its value as a landscape mechanism for sustaining wetland and riparian ecosystems. While the ecological detractors of livestock grazing are well documented in the Intermountain West (Fleischner, 1994; Kauffman and Krueger, 1984; Richter et al., 2020), our results suggest the relationships between ranching and the environment are nuanced, providing opportunities to enhance ecological benefits through sustainable agriculture. The overlap we identify between grass-hay wetlands and historical riparian ecosystems are indicators that provide ample evidence of broad-scale ES. Still, we acknowledge these benefits can be associated with trade-offs stemming from the overallocation of surface water diversions that may impact the integrity of aquatic resources (sensu Alger et al., 2021). As climate change impacts mount, greater awareness of these effects will be necessary to manage water resources effectively for agricultural and ecological benefits.

Cost-effective implantation of public policy incentives for agricultural ES requires a better understanding of potential ecological outcomes (Swinton et al., 2007). Known concentrations of ES in 2.5% of irrigated lands provide targeted conservation opportunities, supporting nearly 60% and 20% of temporary and seasonal wetlands in the Intermountain West. As such, we urge federal and state natural resource agencies to consider the development of volunteer incentive programs targeting the sustainability of ranching and grass-hay production practices linked to the conservation of wetlands and riparian ecosystems. Recent case studies in the Beaverhead and Teton River basins in Montana and Idaho have demonstrated that incentivization of grass-hay flood irrigation can be used as an effective tool to increase groundwater recharge and maintain late summer stream flows (Lonsdale et al., 2020).

We make our grass-hay wetland data publicly available as interactive web-based mapping tools for conservation design. Tool use requires local and regional considerations of specific social, ecological, and economic factors discussed herein to accurately identify risks and opportunities to sustain wetland and riparian resources. We encourage the use of our results to inform conservation solutions through collaborative and proactive decision-making among resource stakeholders.

## CRediT authorship contribution statement

J Patrick Donnelly conceived the analysis. J Patrick Donnelly carried out the analysis. All authors discussed and revised the manuscript.

## Declaration of Competing Interest

The authors declare that they have no known competing financial interests or personal relationships that could have appeared to influence the work reported in this paper.

## Acknowledgments

Views in this manuscript from United States Fish and Wildlife Service authors are their own and do not necessarily represent the agency’s views. Any use of trade, firm, or product names is for descriptive purposes only and does not imply endorsement by the U.S. Government.

## Data availability

All wetland surface water data developed for this study may be accessed as interactive maps from https://4932539.users.earthengine.app/view/wwex. Any further data requests should be directed to the lead author.

## Notes

### Competing Interest Statement

The authors have declared no competing interest.

### Summary of Updates

Figures

## References

Alger, M., Lane, B.A., Neilson, B.T., 2021. Combined influences of irrigation diversions and associated subsurface return flows on river temperature in a semi-arid region. Hydrol. Process. 35. 10.1002/hyp.14283

Allred, B.W., Bestelmeyer, B.T., Boyd, C.S., Brown, C., Davies, K.W., Duniway, M.C., Ellsworth, L.M., Erickson, T.A., Fuhlendorf, S.D., Griffiths, T.V., Jansen, V., Jones, M.O., Karl, J., Knight, A., Maestas, J.D., Maynard, J.J., McCord, S.E., Naugle, D.E., Starns, H.D., Twidwell, D., Uden, D.R., 2021. Improving Landsat predictions of rangeland fractional cover with multitask learning and uncertainty. Methods Ecol. Evol. 12, 841– 849.

[BHWC] Big Hole Watershed Committee, 2023. Big Hole River Drought Management Plan (No. Version 2023). Big Hole Watershed Committee.

Cardille, J.A., Crowley, M.A., Saah, D., Clinton, N.E., 2023. Cloud-Based Remote Sensing with Google Earth Engine: Fundamentals and Applications. Springer Nature.

Cohen, J., 1960. A Coefficient of Agreement for Nominal Scales. Educ. Psychol. Meas. 20, 37–46.

Cowardin, L.M., Carter, F.C., Golet, E.T., 1979. Classification of wetlands and deepwater habitats of the United States. United States Department of the Interior, Fish and Wildlife Service, Washington, DC, USA.

Dieter, C.A., 2018. Water Availability and Use Science Program: Estimated Use of Water in the United States In 2015. U.S. Geological Survey.

Döll, P., Siebert, S., 2002. Global modeling of irrigation water requirements. Water Resour. Res. 38, 8-1 – 8-10.

Donnelly, J.P., Allred, B.W., Perret, D., Silverman, N.L., Tack, J.D., Dreitz, V.J., Maestas, J.D., Naugle, D.E., 2018. Seasonal drought in North America’s sagebrush biome structures dynamic mesic resources for sage-grouse. Ecol. Evol. 8, 12492–12505.

Donnelly, J.P., King, S.L., Knetter, J., Gammonley, J.H., Dreitz, V.J., Grisham, B.A., Nowak, M.C., Collins, D.P., 2021. Migration efficiency sustains connectivity across agroecological networks supporting sandhill crane migration. Ecosphere 12, 1–22.

Downard, R., Endter-Wada, J., 2013. Keeping wetlands wet in the western United States: adaptations to drought in agriculture-dominated human-natural systems. J. Environ. Manage. 131, 394–406.

Dozier, A.Q., Arabi, M., Wostoupal, B.C., Goemans, C.G., Zhang, Y., Paustian, K., 2017. Declining agricultural production in rapidly urbanizing semi-arid regions: policy tradeoffs and sustainability indicators. Environ. Res. Lett. 12, 1–9.

Essaid, H.I., Caldwell, R.R., 2017. Evaluating the impact of irrigation on surface water - groundwater interaction and stream temperature in an agricultural watershed. Sci. Total Environ. 599–600, 581–596.

European Space Agency, 2015. Sentinel-2 Mission Guide [WWW Document]. Sentinel Online. URL https://sentinels.copernicus.eu/web/sentinel/missions/sentinel-2 (accessed 2.26.23).

Ferencz, S.B., Tidwell, V.C., 2022. Physical Controls on Irrigation Return Flow Contributions to Stream Flow in Irrigated Alluvial Valleys. Frontiers in Water 4. 10.3389/frwa.2022.828099

Fleischner, T.L., 1994. Ecological Costs of Livestock Grazing in Western North America. Conserv. Biol. 8, 629–644.

Fleskes, J.P., Yee, J.L., 2007. Waterfowl distribution and abundance during spring migration in southern Oregon and northeastern California. N. Am. Nat. 67, 409–428.

Google, 2022. Google Satellite Tiles [WWW Document]. Google Maps Platform. URL https://developers.google.com/maps/documentation/tile/satellite (accessed 7.26.23).

Gordon, B.L., Paige, G.B., Miller, S.N., Claes, N., Parsekian, A.D., 2020. Field scale quantification indicates potential for variability in return flows from flood irrigation in the high altitude western US. Water Manage. 232, 1–12.

Gorelick, N., Hancher, M., Dixon, M., Ilyushchenko, S., Thau, D., Moore, R., 2017. Google Earth Engine: Planetary-scale geospatial analysis for everyone. Remote Sens. Environ. 202, 18–27.

Gornall, J., Betts, R., Burke, E., Clark, R., Camp, J., Willett, K., Wiltshire, A., 2010. Implications of climate change for agricultural productivity in the early twenty-first century. Philos. Trans. R. Soc. Lond. B Biol. Sci. 365, 2973–2989.

Hansen, Z.K., Libecap, G.D., Lowe, S.E., 2009. Climate Variability and Water Infrastructure: Historical Experience in the Western United States (No. 15558), Working Paper Series. National Bureau of Economic Research. 10.3386/w15558

Hrozencik, R.A., Aillery, M., 2021. Trends in U.S. Irrigated Agriculture: Increasing Resilience Under Water Supply Scarcity (No. 229). U.S. Department of Agriculture, Economic Research Service. 10.2139/ssrn.3996325

Kauffman, B.J., Krueger, W.C., 1984. Livestock impacts on riparian ecosystems and streamside management implications… A review. J. Range Manage. 37, 403–437.

Ketchum, D., Jencso, K., Maneta, M.P., Melton, F., Jones, M.O., Huntington, J., 2020. IrrMapper: A Machine Learning Approach for High Resolution Mapping of Irrigated Agriculture Across the Western U.S. Remote Sensing 12, 2328.

King, M., Altdorff, D., Li, P., Galagedara, L., Holden, J., Unc, A., 2018. Northward shift of the agricultural climate zone under 21st-century global climate change. Scientific Reports 8, 1–10.

Lillesand, T., Kiefer, R.W., Chipman, J., 2015. Remote Sensing and Image Interpretation. John Wiley & Sons.

Lonsdale, W.R., Cross, W.F., Dalby, C.E., Meloy, S.E., Schwend, A.C., 2020. Evaluating Irrigation Efficiency: Toward a Sustainable Water Future for Montana. Montana State University. 10.1111/gwat.12413

McGuire, C.R., Nufio, C.R., Bowers, M.D., Guralnick, R.P., 2012. Elevation-dependent temperature trends in the Rocky Mountain Front Range: changes over a 56- and 20-year record. PLoS One 7, e44370.

[MEA] Millennium Ecosystem Assessment, 2005. Ecosystems and human well-being. Island press Washington, DC.

Melillo, J.M., Richmond, T., Yohe, G.W., 2014. Climate change impacts in the United States: U.S. national climate assessment. U.S. Global Change Research Program.

Miller, M.R., Takekawa, J.Y., Fleskes, J.P., Orthmeyer, D.L., Casazza, M.L., Perry, W.M., 2005. Spring migration of Northern Pintails from California’s Central Valley wintering area tracked with satellite telemetry: routes, timing, and destinations. Can. J. Zool. 83, 1314–1332.

Milly, P.C.D., Dunne, K.A., 2020. Colorado River flow dwindles as warming-driven loss of reflective snow energizes evaporation. Science 367, 1252–1255.

Minckley, T.A., Turner, D.S., Weinstein, S.R., 2013. The relevance of wetland conservation in arid regions: A re-examination of vanishing communities in the American Southwest. J. Arid Environ. 88, 213–221.

[NASS] USDA National Agricultural Statistics Service, 2019. USDA National Agricultural Statistics Service Cropland Data Layer.

[NRCS] USDA Natural Resources Conservation Service, 2010. USDA Common Land Unit [WWW Document]. Common Land Unit (CLU) INFORMATION SHEET. URL https://www.fsa.usda.gov/Internet/FSA_File/clu_2007_infosheetpdf.pdf (accessed 7.26.23).

Patten, D.T., 1998. Riparian ecosytems of semi-arid North America: Diversity and human impacts. Wetlands 18, 498–512.

Pettorelli, N., Vik, J.O., Mysterud, A., Gaillard, J.-M., Tucker, C.J., Stenseth, N.C., 2005. Using the satellite-derived NDVI to assess ecological responses to environmental change. Trends Ecol. Evol. 20, 503–510.

Power, A.G., 2010. Ecosystem services and agriculture: tradeoffs and synergies. Philos. Trans. R. Soc. Lond. B Biol. Sci. 365, 2959–2971.

QGIS Development Team, 2020. QGIS. Open Source Geospatial Foundation Project.

Rajagopalan, B., Lall, U., 1998. Interannual variability in western US precipitation. J. Hydrol. 210, 51–67.

R Core Team, 2019. R: A Language and Environment for Statistical Computing. R Foundation for Statistical Computing, Vienna, Austria.

Richter, B.D., Bartak, D., Caldwell, P., Davis, K.F., Debaere, P., Hoekstra, A.Y., Li, T., Marston, L., McManamay, R., Mekonnen, M.M., Ruddell, B.L., Rushforth, R.R., Troy, T.J., 2020. Water scarcity and fish imperilment driven by beef production. Nature Sustainability 3, 319–328.

Richter, B.D., Brown, J.D., DiBenedetto, R., Gorsky, A., Keenan, E., Madray, C., Morris, M., Rowell, D., Ryu, S., 2017. Opportunities for saving and reallocating agricultural water to alleviate water scarcity. Water Policy 19, 886–907.

RStudio Team, 2019. RStudio: Integrated Development Environment for R.

Ryder, H.B., 2021. Wall Street Eyes Billions in the Colorado’s Water. New Your Times.

Scott, C.A., Vicuña, S., Blanco Gutiérrez, I., Meza, F., Varela Ortega, C., 2014. Irrigation efficiency and water-policy implications for river-basin resilience. Hydrol. Earth Syst. Sci. 18, 1339–1348.

Silverman, N.L., Allred, B.W., Donnelly, J.P., Chapman, T.B., Maestas, J.D., Wheaton, J.M., White, J., Naugle, D.E., 2019. Low-tech riparian and wet meadow restoration increases vegetation productivity and resilience across semiarid rangelands. Restor. Ecol. 27, 269–278.

Swinton, S.M., Lupi, F., Robertson, G.P., Hamilton, S.K., 2007. Ecosystem services and agriculture: Cultivating agricultural ecosystems for diverse benefits. Ecol. Econ. 64, 245–252.

Talbert, C.B., Knight, R.L., Mitchell, J.E., 2007. Private Ranchlands and Public Land Grazing in the Southern Rocky Mountains: Why the private land matters when we think about public lands grazing. Rangelands 29, 5–8.

Thorvaldson, J., Pritchett, J.G., 2006. Economic impact analysis of reduced irrigated acreage in four river basins in Colorado. Colorado Water Resources Research Institute. https://watercenter.colostate.edu/wp-content/uploads/sites/33/2020/03/CR207.pdf

Tiner, R.W., 2003. Estimated extent of geographically isolated wetlands in selected areas of the United States. Wetlands 23, 636.

Tracy, B., Bast, A., Spinder, C., 2023. New York investors snapping up Colorado River water rights, betting big on an increasingly scarce resource. CBS News.

[USGS] United States Department of Interior, Geological Survey, 2012. National Elevation Dataset 1/3 arc-second.

[USGS] United States Geological Survey, Gap Analysis Project (GAP), 2018. Protected Areas Database of the United States (PAD-US). 10.5066/P955KPL

Wallander, S., Hand, M.S., 2011. Measuring the impact of the environmental quality incentives program (EQIP) on irrigation efficiency and water conservation. Agricultural and Applied Economics Association.

Welsh, L., Endter-Wada, J., Downard, R., Kettenring, K., 2013. Developing Adaptive Capacity to Droughts: the Rationality of Locality. Ecol. Soc. 18. 10.5751/ES-05484-180207

Welsh, L.W., Endter-Wada, J., 2017. Piping water from rural counties to fuel growth in Las Vegas, Nevada: Water transfer risks in the arid USA West. Water Alternatives 10, 420.

Wickham, H., Averick, M., Bryan, J., Chang, W., McGowan, L.D., François, R., Grolemund, G., Hayes, A., Henry, L., Hester, J., Kuhn, M., Pedersen, T.L., Miller, E., Bache, S.M., Müller, K., Ooms, J., Robinson, D., Seidel, D.P., Spinu, V., Takahashi, K., Vaughan, D., Wilke, C., Woo, K., Yutani, H., 2019. Welcome to the tidyverse. Journal of Open Source Software 4, 1686.

Wiken, E., Jiménez Nava, F., Griffith, G., 2011. North American Terrestrial Ecoregions—Level III. Commission for Environmental Cooperation, Montreal, Canada.

Zhuang, W., 2016. Eco-environmental impact of inter-basin water transfer projects: a review. Environ. Sci. Pollut. Res. 23, 12867–12879.

