## Supplementary material for "Beneficial ‘inefficiencies’ of western ranching: Flood-irrigated hay production sustains wetland systems by mimicking historic hydrologic processes": sup

1. Remote Sensing Methods

1.1 Surface water modeling

Following an approach outlined by Donnelly et al. (2021), monthly surface water hydrology was measured from Landsat multispectral imagery using constrained spectral mixture analysis (SMA, Adams and Gillespie, 2006) that allowed proportional estimations of water contained within a continuous 30 x 30 m pixel grid (Halabisky et al., 2016; Jin et al., 2017). This approach accurately accounts for surface water area/extent when detectability is reduced due to interspersion of emergent vegetation, shallow, or turbid water (DeVries et al., 2017), characteristics common to flood-irrigation and other water bodies in the Intermountain West. Pixels were considered fully inundated when water was present because emergent vegetation or high turbidity could partially mask pixel areas covered with water (Donnelly et al., 2019). Pixels containing <15% surface water were omitted from summaries to minimize the overestimation of surface water area. Satellite data containing clouds, cloud shadows, snow, and ice were masked using the Landsat CFMask band (Foga et al., 2017). All unmasked pixels in Landsat 30 m visible, near-infrared, and short-wave infrared bands were incorporated into the SMA except for the Landsat 8 coastal aerosol band.

Training data for SMA were extracted from satellite imagery as spectral endmembers unique to individual images classified. Training site locations represented homogeneous land cover types mapped as water, wetland vegetation, upland, and bare soil. Spectral endmembers for water were collected using image masks generated from 99th percentile normalized difference water index values (McFeeters, 1996). Mask extents coincided with large deep water lakes/reservoirs proximal to wetland sites. A similar masking approach was applied to collect wetland vegetation endmembers using normalized difference vegetation indices (Box et al., 1989). Sampling was constrained to sites coincident with flooded wetlands and were representative of associated plant phenology. Spectral mixture analysis requires minimal training data (Adams and Gillespie, 2006), which allowed upland and bare soil endmembers to be generated from a small number of static plots within the study area (n = 2; 0.5-1 km^2^). Upland plots were associated with shrub-steppe rangelands. Bare soil plots were coincident with dry lake basins in areas of surface mineral deposits. Plot locations were identified using high-resolution (≤ 1-meter) multispectral satellite imagery or field survey.

Using digital aerial orthophoto interpretation, wetland delineations were classified into functional groups to identify specific ecologic and land-use characteristics associated with surface water hydrology. Classes included lake, reservoir, swedge pond, and large rivers. This made it possible to stratify wetland data by removing extraneous or large deep water (e.g., lake and reservoir) features that would have biased results. The accuracy of surface water area was estimated to be 93-98% by comparison to previous work and similar methods used by Donnelly et al. (2019) that overlapped a quarter of our study site. Accuracy was comparable to similar time‐series wetland inundation studies using Landsat data (Jin, Huiran Huang, Chengquan Lang, Megan W Yeo, In-Young Stehman, Stephen V, 2017).

1.2 Majority Filter Application

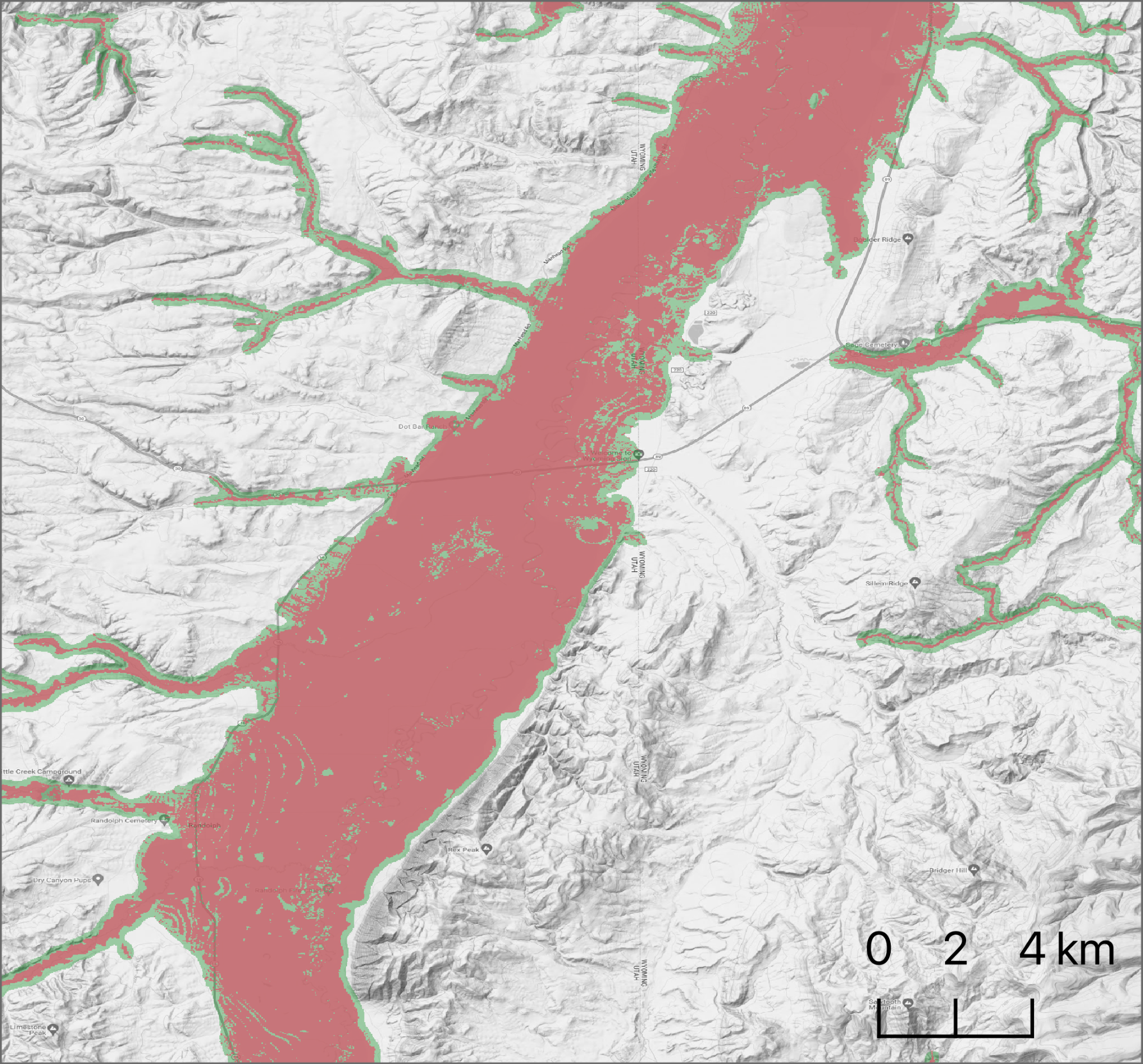

Fig. S1. Example comparison of aggregated riparian ecosystem classes from LANDFIRE BioPhysical Settings layer extent pre (rust) and post (rust + green) application of majority filter. Results normalized minor areas of non-riparian pixels for a more heterogeneous representation of historic riparian ecosystem extent.

Figures S2-S10 depict state-level summaries of mean monthly wetland surface water by hydroperiod class (temporary, seasonal, and semi-permanent) from 2013-22. Measures shown for wetlands and wetlands supported by flood-irrigated grass-hay production. Only portions of states included in the Intermountain West study are included (See Fig. 1)

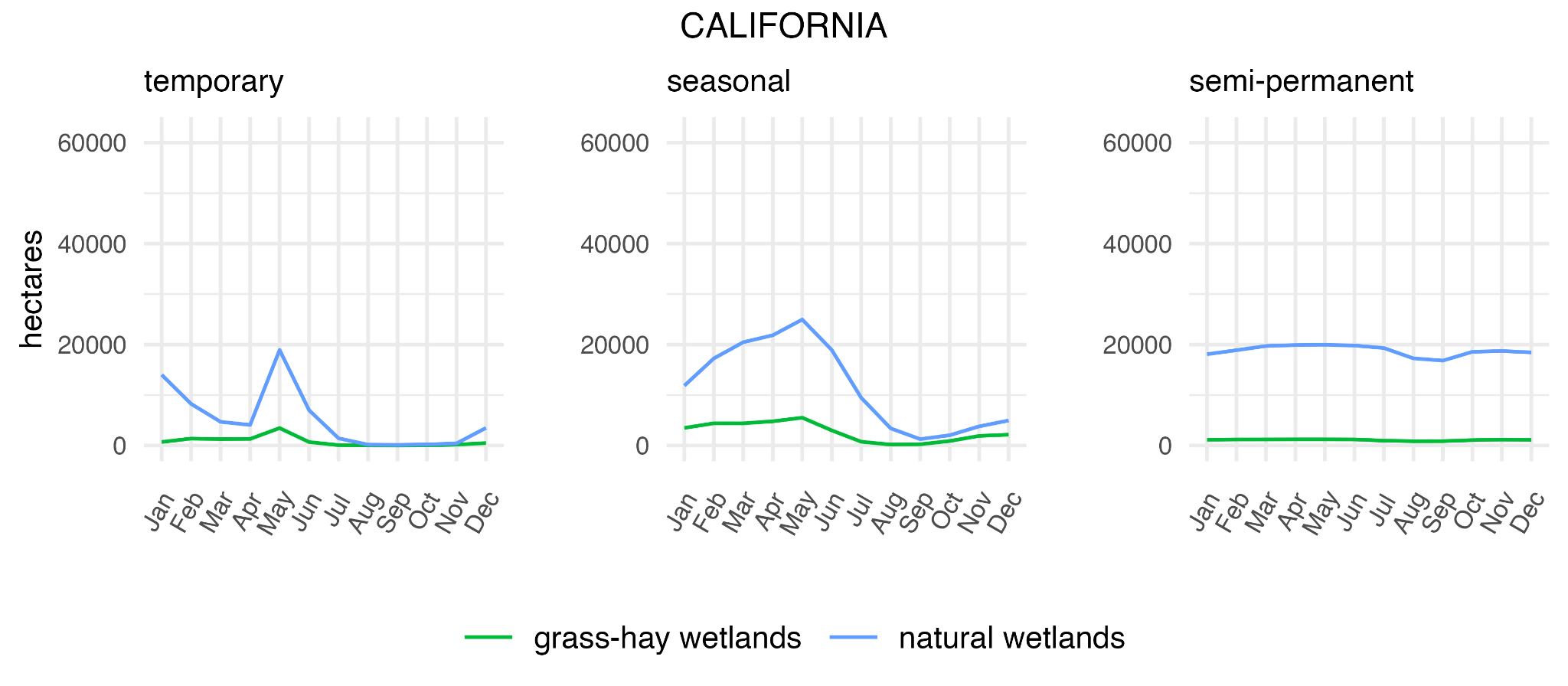

Fig. S2. California mean monthly wetland surface water by hydroperiod class (temporary, seasonal, and semi-permanent) from 2013-22. Measures shown for wetlands and wetlands supported by flood-irrigated grass-hay production.

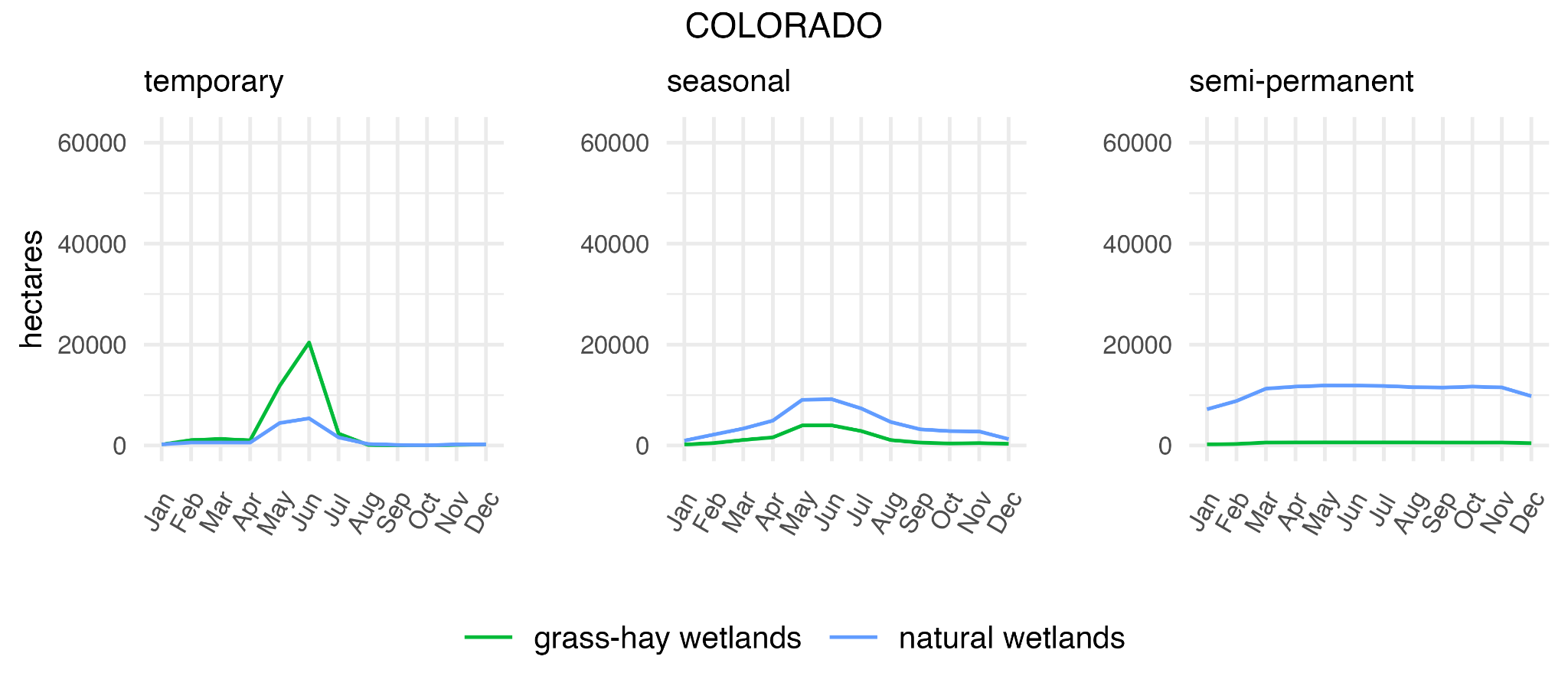

Fig. S3. Colorado mean monthly wetland surface water by hydroperiod class (temporary, seasonal, and semi-permanent) from 2013-22. Measures shown for wetlands and wetlands supported by flood-irrigated grass-hay production.

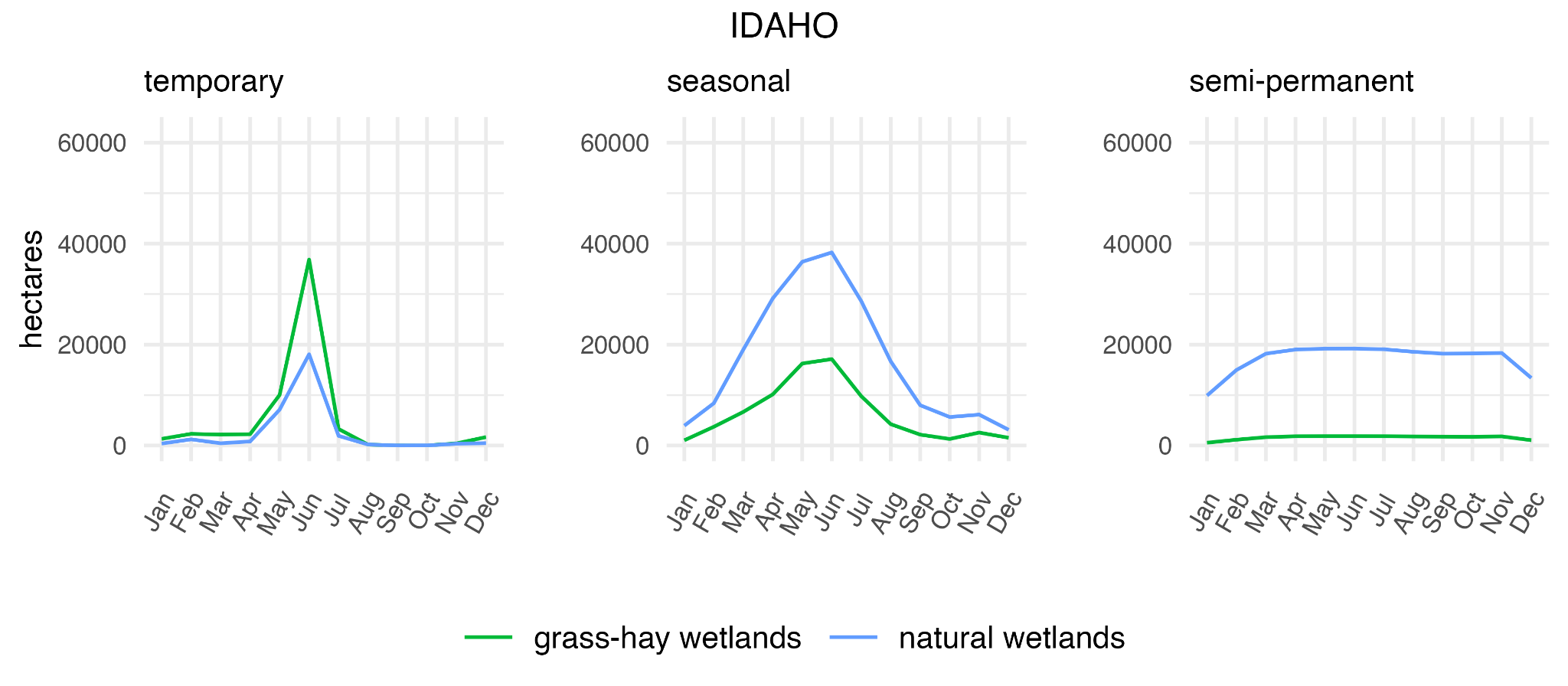

Fig. S4. Idaho mean monthly wetland surface water by hydroperiod class (temporary, seasonal, and semi-permanent) from 2013-22. Measures shown for wetlands and wetlands supported by flood-irrigated grass-hay production.

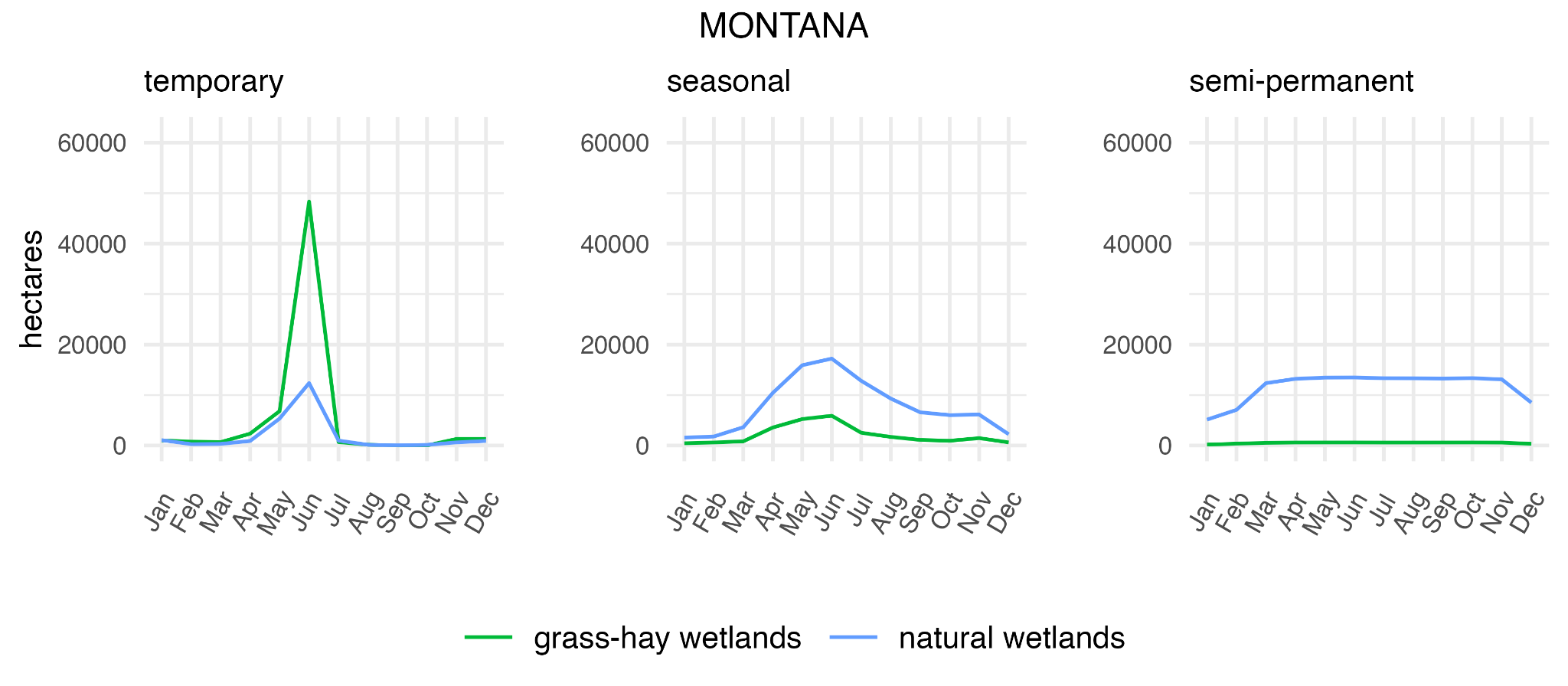

Fig. S5. Montana mean monthly wetland surface water by hydroperiod class (temporary, seasonal, and semi-permanent) from 2013-22. Measures shown for wetlands and wetlands supported by flood-irrigated grass-hay production.

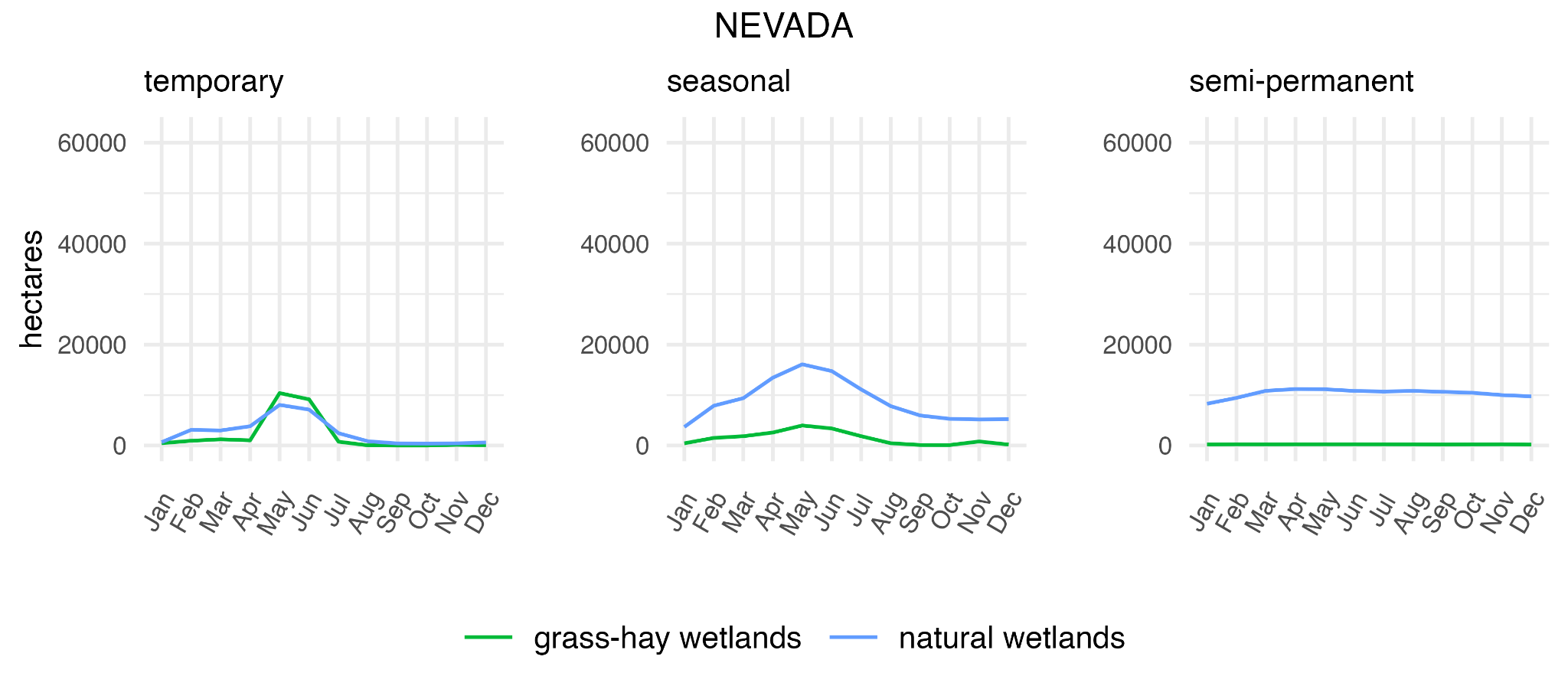

Fig. S6. Nevada mean monthly wetland surface water by hydroperiod class (temporary, seasonal, and semi-permanent) from 2013-22. Measures shown for wetlands and wetlands supported by flood-irrigated grass-hay production.

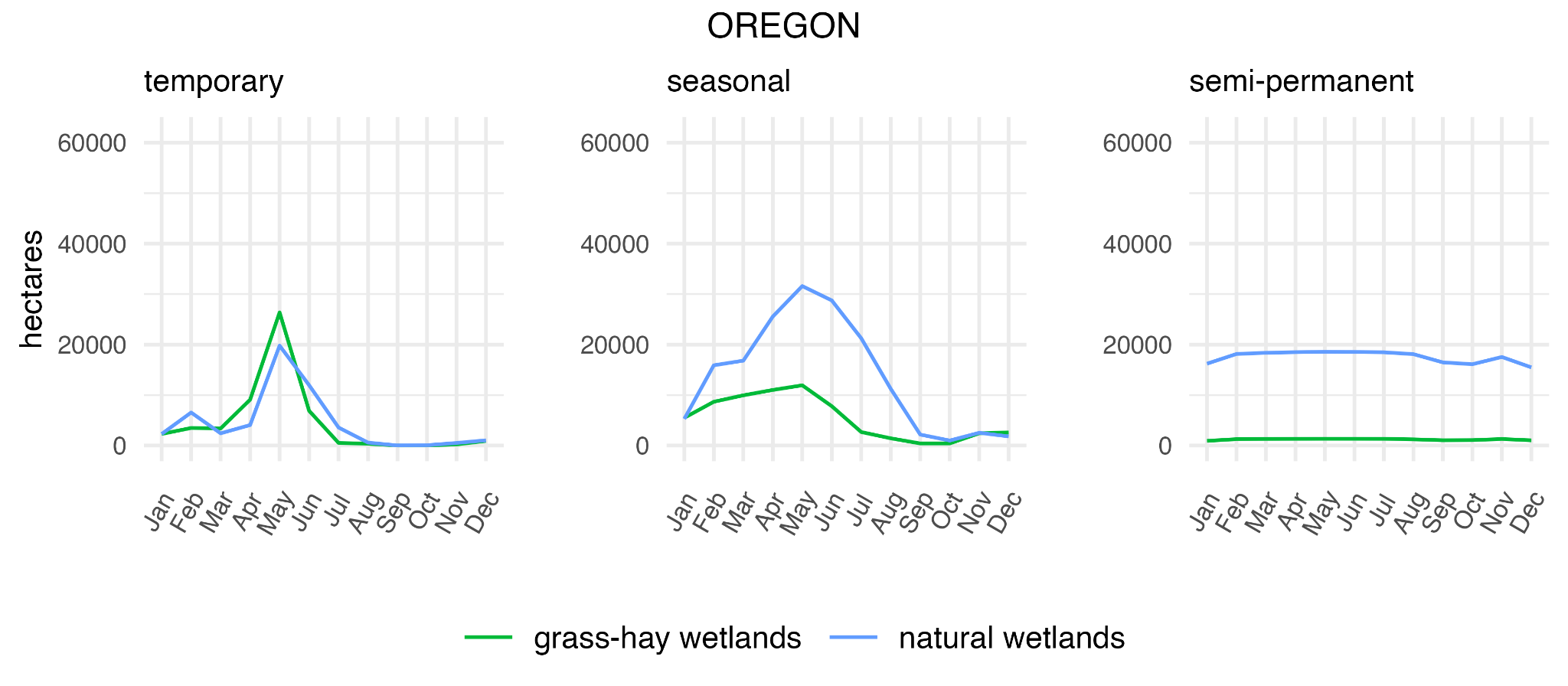

Fig. S7. Oregon mean monthly wetland surface water by hydroperiod class (temporary, seasonal, and semi-permanent) from 2013-22. Measures shown for wetlands and wetlands supported by flood-irrigated grass-hay production.

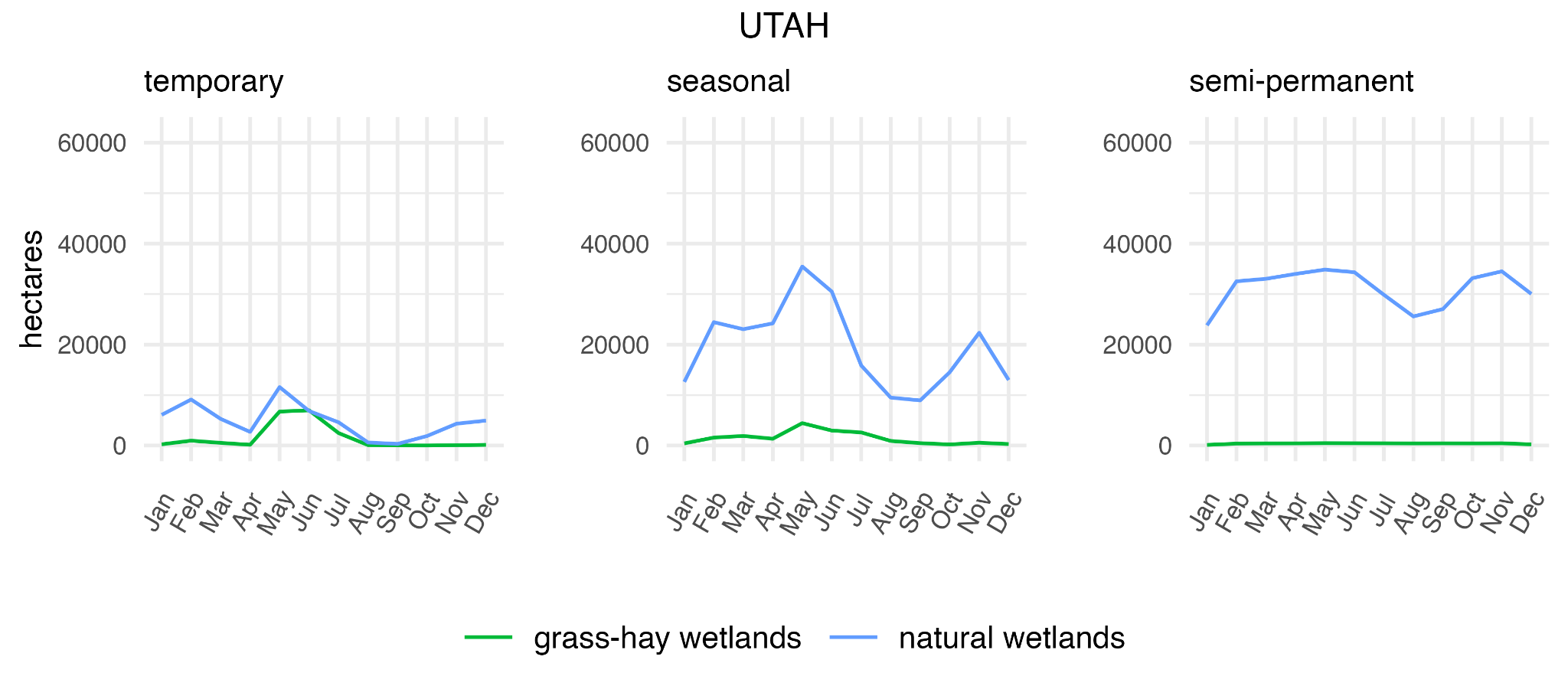

Fig. S8. California mean monthly wetland surface water by hydroperiod class (temporary, seasonal, and semi-permanent) from 2013-22. Measures shown for wetlands and wetlands supported by flood-irrigated grass-hay production.

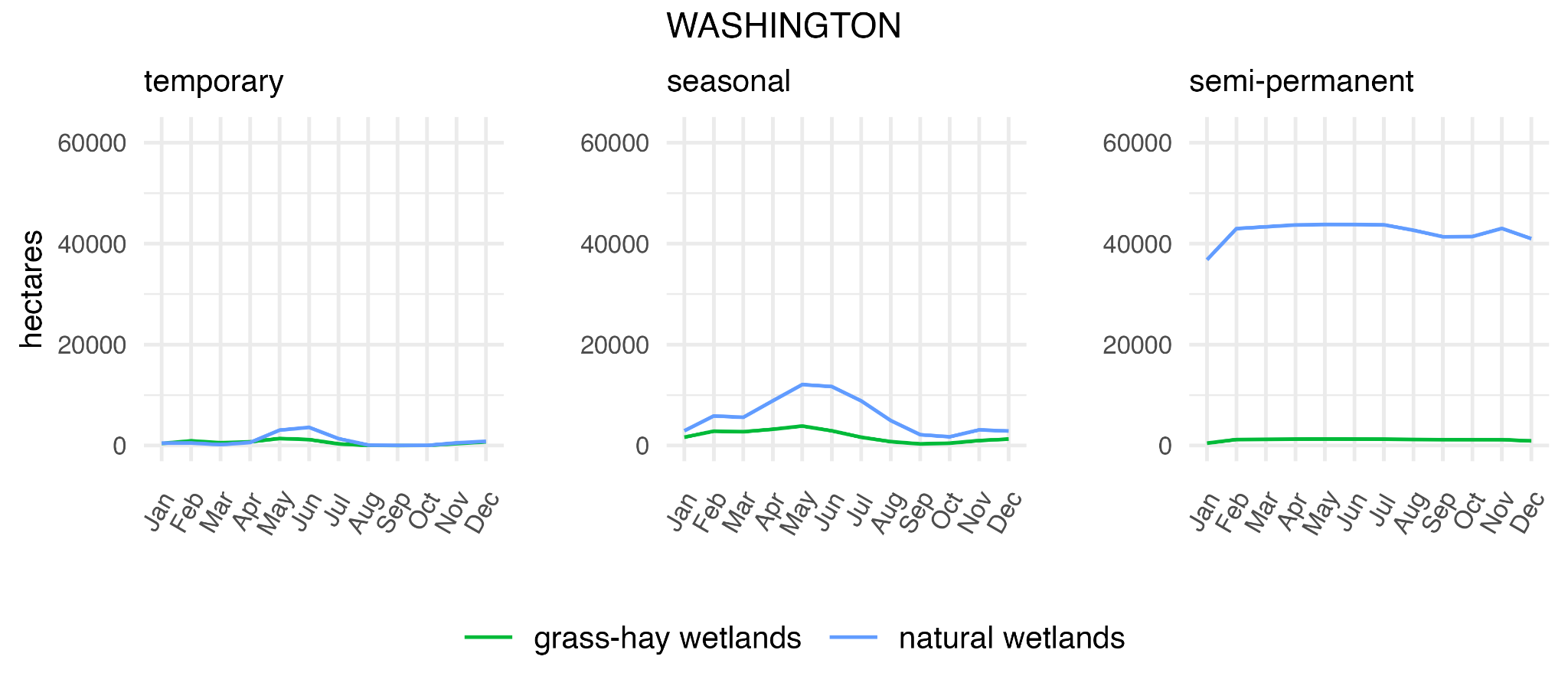

Fig. S9. Washington mean monthly wetland surface water by hydroperiod class (temporary, seasonal, and semi-permanent) from 2013-22. Measures shown for wetlands and wetlands supported by flood-irrigated grass-hay production.

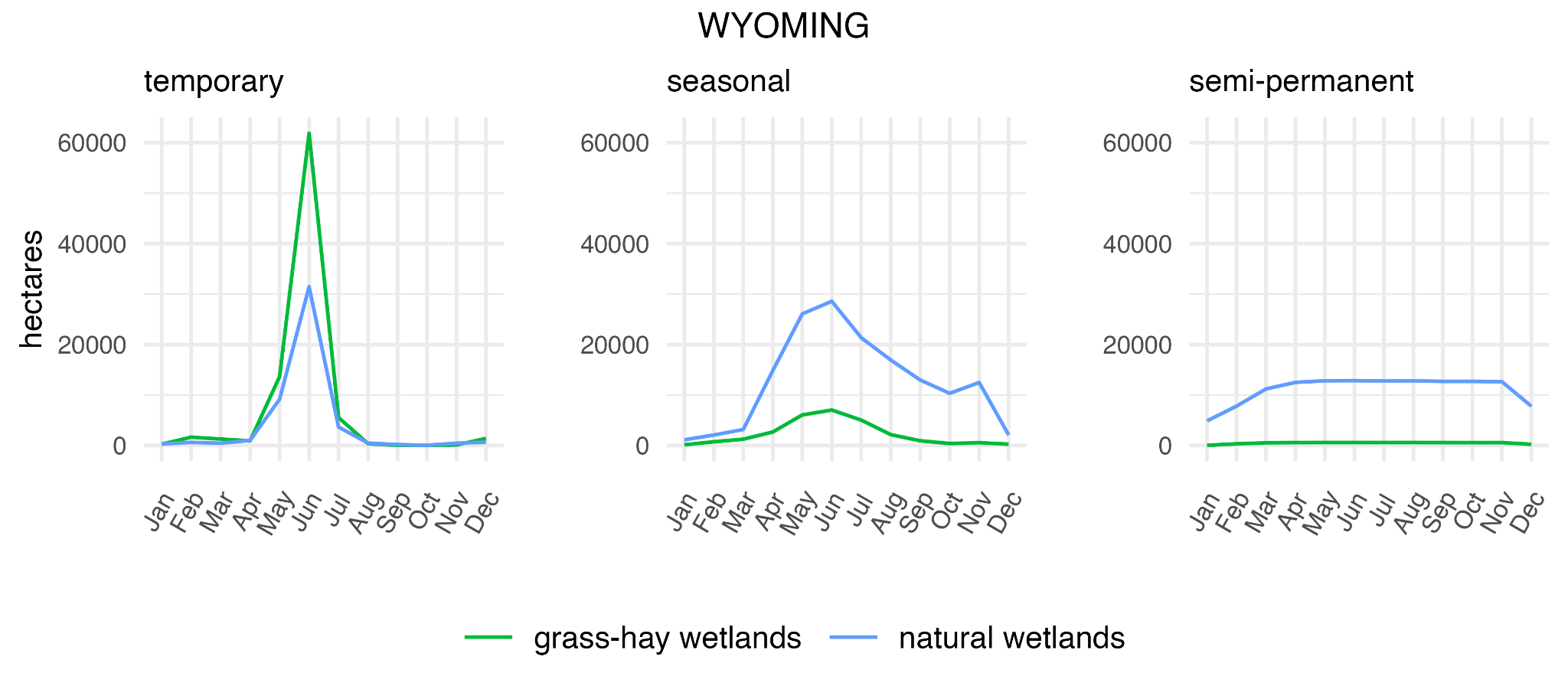

Fig. S10. Wyoming mean monthly wetland surface water by hydroperiod class (temporary, seasonal, and semi-permanent) from 2013-22. Measures shown for wetlands and wetlands supported by flood-irrigated grass-hay production.

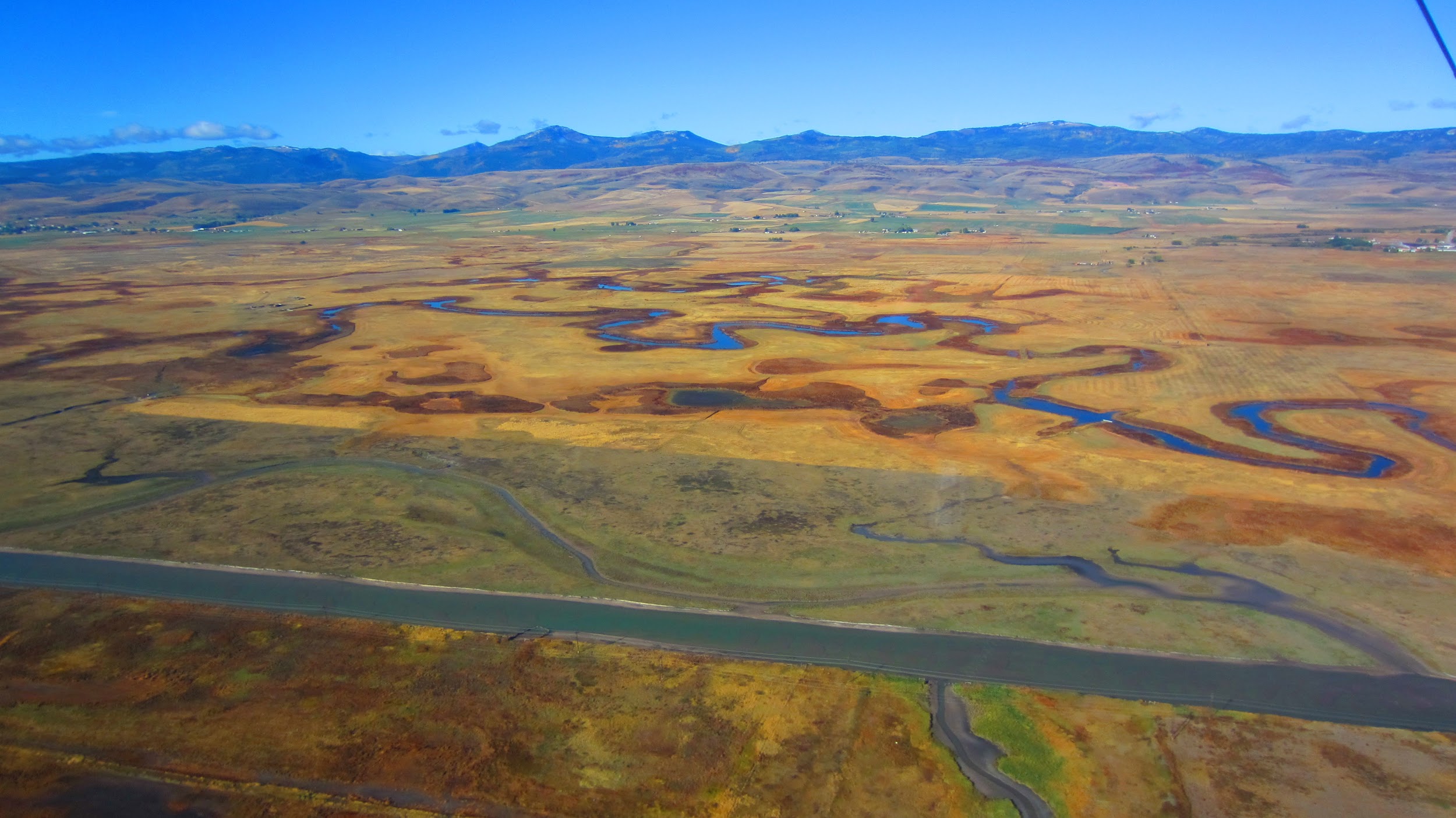

Figure S11. Example of historic river channels supporting semi-permanent wetlands as part of flood-irrigated grass-hay production in the Upper Bear River floodplain, Utah. Lower topographic relief of abandoned channels concentrates surplus surface water.

References

Adams, J.B., Gillespie, A.R., 2006. Remote Sensing of Landscapes with Spectral Images: A Physical Modeling Approach. Cambridge University Press, Cambridge, UK.

Box, E.O., Holben, B.N., Kalb, V., 1989. Accuracy of the AVHRR vegetation index as a predictor of biomass, primary productivity and net CO2 flux. Vegetatio 80, 71–89.

DeVries, B., Huang, C., Lang, M.W., Jones, J.W., Huang, W., Creed, I.F., Carroll, M.L., 2017. Automated Quantification of Surface Water Inundation in Wetlands Using Optical Satellite Imagery. Remote Sensing 9, 807.

Donnelly, J.P., King, S.L., Knetter, J., Gammonley, J.H., Dreitz, V.J., Grisham, B.A., Nowak, M.C., Collins, D.P., 2021. Migration efficiency sustains connectivity across agroecological networks supporting sandhill crane migration. Ecosphere 12. https://doi.org/[10.1002/ecs2.3543](http://dx.doi.org/10.1002/ecs2.3543)

Donnelly, J.P., Naugle, D.E., Collins, D.P., Dugger, B.D., Allred, B.W., Tack, J.D., Dreitz, V.J., 2019. Synchronizing conservation to seasonal wetland hydrology and waterbird migration in semi‐arid landscapes. Ecosphere 10, 459.

Foga, S., Scaramuzza, P.L., Guo, S., Zhu, Z., Dilley, R.D., Beckmann, T., Schmidt, G.L., Dwyer, J.L., Joseph Hughes, M., Laue, B., 2017. Cloud detection algorithm comparison and validation for operational Landsat data products. Remote Sens. Environ. 194, 379–390.

Halabisky, M., Moskal, L.M., Gillespie, A., Hannam, M., 2016. Reconstructing semi-arid wetland surface water dynamics through spectral mixture analysis of a time series of Landsat satellite images (1984–2011). Remote Sens. Environ. 177, 171–183.

Jin, Huiran Huang, Chengquan Lang, Megan W Yeo, In-Young Stehman, Stephen V, 2017. Monitoring of wetland inundation dynamics in the Delmarva Peninsula using Landsat time-series imagery from 1985 to 2011. Remote Sens. Environ. 190, 26–41.

McFeeters, S.K., 1996. The use of the Normalized Difference Water Index (NDWI) in the delineation of open water features. Int. J. Remote Sens. 17, 1425–1432.
